# Structural and Binding Properties of Dps – A Nucleoid Associated Protein

**DOI:** 10.1101/2025.11.14.688440

**Authors:** Matty Gaines, Daniel Parrell, Kaylee Jo Rajek, Steven Garvis, Bryan Sibert, Elizabeth R. Wright

## Abstract

Prokaryotic chromosomal DNA is compacted and protected by nucleoid-associated proteins (NAPs). Among these, the DNA-binding proteins from starved cells (Dps) combine ferroxidase activity with nonspecific DNA binding to form dodecameric assemblies that condense and safeguard the genome under stress. Here, we present an integrated structural and biophysical analysis of *Escherichia coli* (*E. coli*) Dps across a pH range from 3 to 11, combining (i) single-particle cryo-EM reconstructions at ∼1.75 Å resolution, (ii) adaptive Poisson-Boltzmann electrostatic modelling, (iii) electrophoretic mobility shift assays (EMSAs) on a 448 bp DNA fragment, and (iv) cryo-electron tomography (cryo-ET) of Dps-DNA assemblies. Through this analysis, we show that (1) the canonical ferritin-like fold is maintained at all pH values; (2) surface electrostatics shift from highly positive to net negative as the pH increases, modulating DNA affinity (*EC*_50_ values of 73.4 nM at the lowest measurement of pH 5, to 815.5 nM at the highest measurement of pH 11); and (3) short-fragment Dps-DNA complexes form amorphous ∼50 nm globular or tubular complexes rather than lattice co-crystals observed on kilobase-length DNA scaffolds. The findings reveal a possible two-stage assembly model, in which protonation-driven nucleation occurs via the Dps N-terminal lysine residues, followed by lattice ordering on long DNA. Our results further define Dps as a pH-responsive nucleoid compaction factor in bacteria.

## Introduction

In prokaryotes, the organization, compaction, and protection of chromosomal DNA are orchestrated by nucleoid-associated proteins (NAPs), a diverse class of architectural and regulatory elements that fulfill roles analogous to eukaryotic histones [1, 2]. NAPs such as Fis, HU, IHF, and H-NS shape the bacterial nucleoid and broadly influence gene expression [3, 4]. Among these, the DNA-binding protein from starved cells (Dps) stands out for its structural stability and functional duality, as it acts both as a nonspecific DNA-binding protein and a ferritin-like ferroxidase enzyme [5, 6]. Dps oligomerizes into highly symmetric, dodecameric cages that function to compact bacterial DNA during stress, the stationary phase, and nutrient limitation [6–9]. This process not only condenses the nucleoid but also protects DNA from oxidative damage by sequestering iron and detoxifying reactive oxygen species (ROS) [7, 10].

During logarithmic growth, *Escherichia coli* Dps is expressed at a minimal level, with ∼6,000 molecules per cell. However, its expression can increase by more than 30-fold during the stationary phase and/or oxidative stress to approximately 180,000 molecules per cell [6, 11, 12], underscoring its vital role in genome protection. This remarkable abundance of Dps under stress has motivated many structural studies to clarify how its architecture supports both DNA protection and iron sequestration. Dps plays a more specific role in protecting pathogenic species, such as *E. coli*, particularly enterohemorrhagic *E. coli* (EHEC) O157:H7, by binding to and shielding DNA from acid-induced damage in the host stomach [13, 14].

In 1998, using X-ray crystallography, Grant et al. solved the structure of the apo form of *E. coli* Dps dodecamer at a resolution of 1.6 Å (PDB code: 1DPS) [5], revealing a compact tetrahedral architecture composed of 12 subunits, each forming a four-helix bundle. Flexible, lysine-rich N-terminal tails extended from the surface and were implicated in DNA binding [5]. Subsequent structures, including *Deinococcus radiodurans* Dps-1 [15] (PDB code: 2F7N), resolved at 2.0 Å, revealed conserved ferroxidase centers and additional iron channels, highlighting subtle functional divergence among Dps homologs.

More recent advances in structural biology, particularly cryo-electron microscopy (cryo-EM) and cryo-electron tomography (cryo-ET), have significantly expanded our understanding of Dps behavior in solution and DNA-bound states [16]. Cryo-ET studies of *E. coli* Dps co-crystals on long DNA scaffolds revealed the formation of multiple polymorphic lattices, including triclinic and cubic networks, under different buffer and ionic conditions [17, 18]. Similarly, tomograms of Dps–DNA complexes have revealed morphological heterogeneity under both *in vitro* and *in vivo* conditions, ranging from compact globules to extended filaments, suggesting that DNA length and topology may influence the formation of higher-order assemblies [9, 17–20]. Single-particle cryo-EM studies of Dps homologs further support the model that the Dps–DNA interaction is mediated through the N-terminal tails and is sensitive to the charge state and ionic environment [21, 22]. *Mycobacterium smegmatis* Dps2 was recently imaged in complex with DNA at near-atomic resolution, revealing that its N-terminal residues play a central role in nucleoprotein complex formation [23]. Complementary single-molecule experiments have demonstrated that Dps serves as a bridging factor between DNA segments, facilitating the condensation of DNA into biomolecular assemblies reminiscent of phase-separated condensates [24]. These findings highlight Dps functional plasticity in both local DNA compaction and larger-scale nucleoid organization.

Despite these significant contributions, essential questions remain unanswered. While long DNA substrates (>3 kb) support the formation of a crystalline lattice [5, 8, 17, 25], it is unclear how Dps organizes shorter DNA fragments, which may more closely reflect the constrained domains present within the bacterial nucleoid [7, 26]. Furthermore, the influence of pH, an environmental factor that varies dramatically under stress and starvation, on Dps structure, electrostatic properties, and DNA binding remains incompletely understood [15, 21, 27, 28]. Although it is known that protonation and deprotonation of surface residues, particularly lysine and glutamate, can modulate DNA affinity [22, 29], no study to date has examined the structural and functional behavior of Dps across the full physiological and extreme pH spectrum.

Here, we present high-resolution, pH-resolved structural and functional analyses of *E. coli* Dps, using multiple complementary approaches: (1) single-particle cryo-EM reconstructions at pH 3, 5, 7, 9, and 11 [23]; (2) adaptive Poisson–Boltzmann electrostatic modeling to visualize surface charge distributions; (3) electrophoretic mobility shift assays (EMSAs) to quantify DNA-binding affinities on a 448 bp linear DNA fragment (138.7 kDa); and (4) cryo-ET segmentation of Dps–DNA complexes at critical pH points. Across this wide pH range, we find that the Dps dodecameric ferritin-like core remains structurally stable [20]. At the same time, DNA affinity is strongly pH-dependent, with maximal binding occurring near pH 5 and dramatic reductions at alkaline pH levels. Furthermore, we demonstrate that short DNA fragments facilitate the formation of amorphous Dps–DNA globular complexes, which represent a distinct assembly regime and differ from the crystalline co-assemblies observed with longer DNA. Together, these results provide new insight into the pH-regulated binding behavior of Dps, its structural robustness, and its assembly mechanisms on sub-genomic DNA scales. By integrating structural biology with biophysical assays, this work provides a comprehensive mechanistic model of Dps function as a pH-responsive NAP and a nucleoid compaction factor in bacteria.

## Results

### Cryo-EM Structure of Dps Across pH Values

We initially aimed to determine whether pH extremes induced any structural changes in the *E. coli* Dps protein using single particle cryo-electron microscopy (cryo-EM). Purified Dps protein was pH-shifted to values of 3, 5, 7, 9, and 11 before being plunge-frozen onto cryo-EM grids. Cryo-EM data were collected for Dps at five different pH conditions, and representative raw micrographs from each Dps dataset are shown in Supplementary Figure 1.

Previous work focused on solving the *E. coli* Dps protein structure had already confirmed its dodecamer composition and symmetry value [5]; however, to our knowledge, no prior structural work has been conducted at pH values other than ∼7. To confirm whether structural changes might occur that would break the known symmetry pattern, we determined 3D maps for each dataset without applying symmetry. Each Dps map yielded a resolution of less than 2 Å (Supplementary Figure 2). The 3D structures show that Dps exhibits tetrahedral (T) type symmetry at each pH value, with no gross differences in the structural organization of the oligomers, as observed in previously solved Dps structures at pH 7 [5]. Imposing T symmetry and several rounds of particle-based CTF refinement, (per-particle) motion correction, as well as 3D classification, ultimately lead to maps with values of 1.72 Å, 1.76 Å, 1.86 Å, 1.72 Å, and 1.81 Å at pH 3, 5, 7, 9, and 11, respectively (Figure 1, Supplementary Figure 3, Table 1). Local resolution estimates indicated that the maps were best resolved within the middle alpha helix of each subunit, reaching the Nyquist limit of 1.66 Å. The inner and outer surfaces were less resolved, reaching approximately 1.95 Å in the least-resolved map at pH 7 (Supplementary Figure 4). Each of the final maps showed that the Dps protein consists of 12 identical subunits (Figure 1). To determine the similarity between the cryo-EM maps, the ‘Fit to map’ function in UCSF Chimera [30] was used to calculate cross-correlation scores for each map relative to the pH 7 density. The results showed that each map had over 99% similarity, with the lowest value at pH 9 (99.14%) and the highest at pH 5 (99.83%) (Supplementary Figure 5, Table 2). Notably, even at pH 3 and pH 11, where partial unfolding or dissociation might be expected, the core structure of each subunit remained intact, and the dodecamer retained T symmetry. Due to the extremely high similarity between the maps, the model built from the pH 7 map was docked into the other four structures. Figure 2 highlights the high-quality fit between model and map (Q score) across the pH range. It is remarkable that even at pH 3 and pH 11, conditions at which many proteins would irreversibly unfold, Dps retained its quaternary structure long enough to be vitrified and imaged at high resolution. We noted that the cryo-EM sample at pH 11 had fewer intact particles overall, yielding ∼50% of the particle count of the other conditions, and more broken protein background in micrographs, suggesting that some Dps dodecamers dissociated on the grid. At pH 3, the particle images appeared a bit “fuzzier” and occasionally irregular in shape, possibly due to aggregation or partial unfolding, which will be discussed when we examine 2D lattice formation.

**Figure 1.**
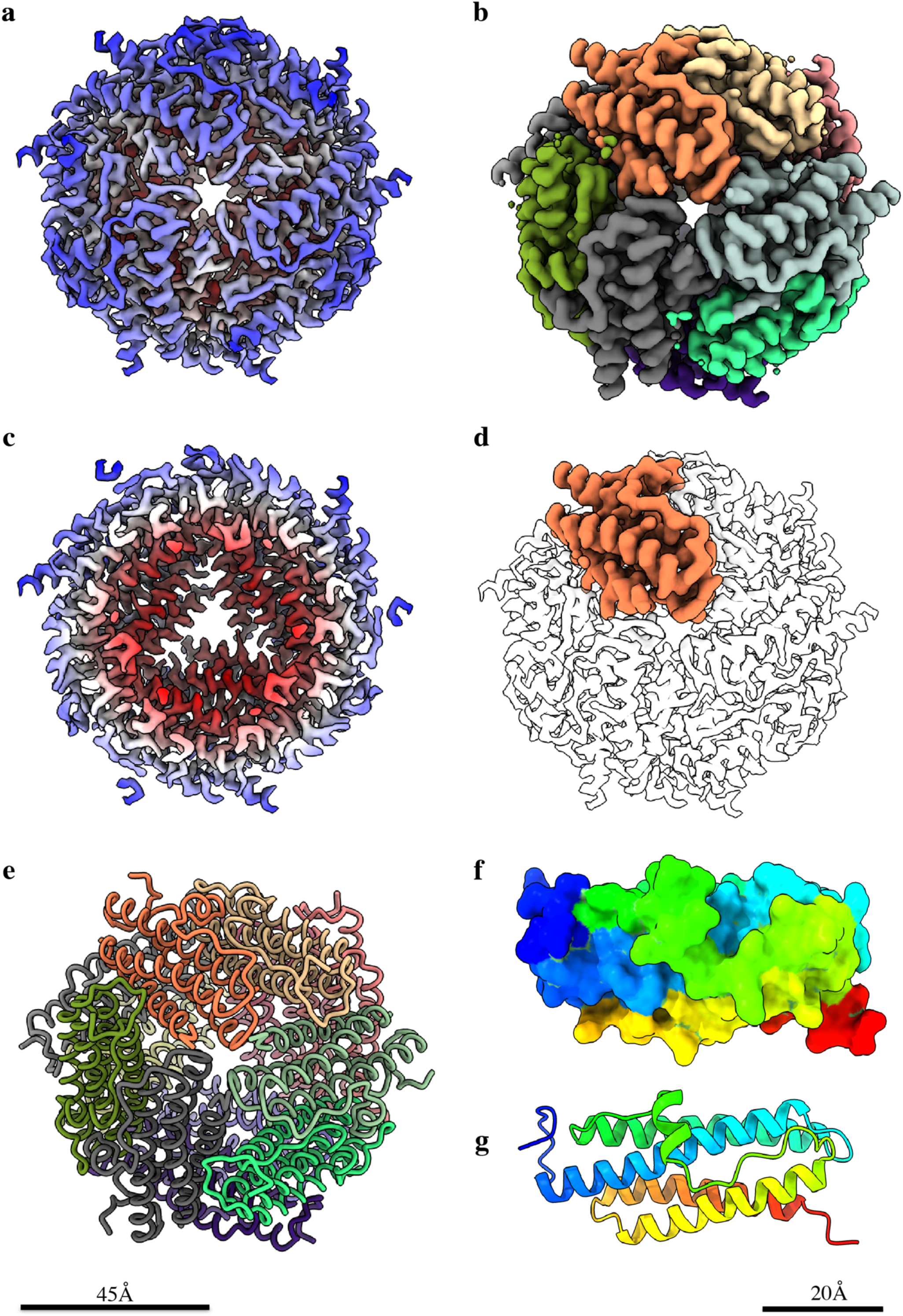
Protein reconstruction and atomic model of the pH 7 *E.coli* Dps. Surface (**a, b**) and cross-sectional (**c, d**) views of the cryoEM map of pH 7 Dps. In **a, c,** the maps are colored by cylinder radius (hollow core surrounded by α-helices, red; α-helical outer shell, in shades of blue). In **b** and **d,** the map is colored by subunit, where **b** shows all 12 subunits, and **d** highlights just one Dps monomer in the electron density map. (**e**) shows the atomic model of Dps with proteins visualized in licorice view. Additionally, one Dps monomer is shown in rainbow color coding, from N to C terminus, in surface (**f**) and ribbon (**g**) formats. Observing (**g**) allows visualization of the individual secondary structure components of the DPS monomer, with the N termini shown in dark blue, alpha-helix 1 (α1) in blue/cyan, and alpha-helix 2 (α2) in cyan/green. The flexible linker region with a small α-helix coil in the middle is shown in green, α-3 in yellow, α-4 in orange/red, and ultimately the C-terminal in red. Scale bars for (**a** – **e**) 45 Å, (**f** and **g**) 20 Å.

**Figure 2.**
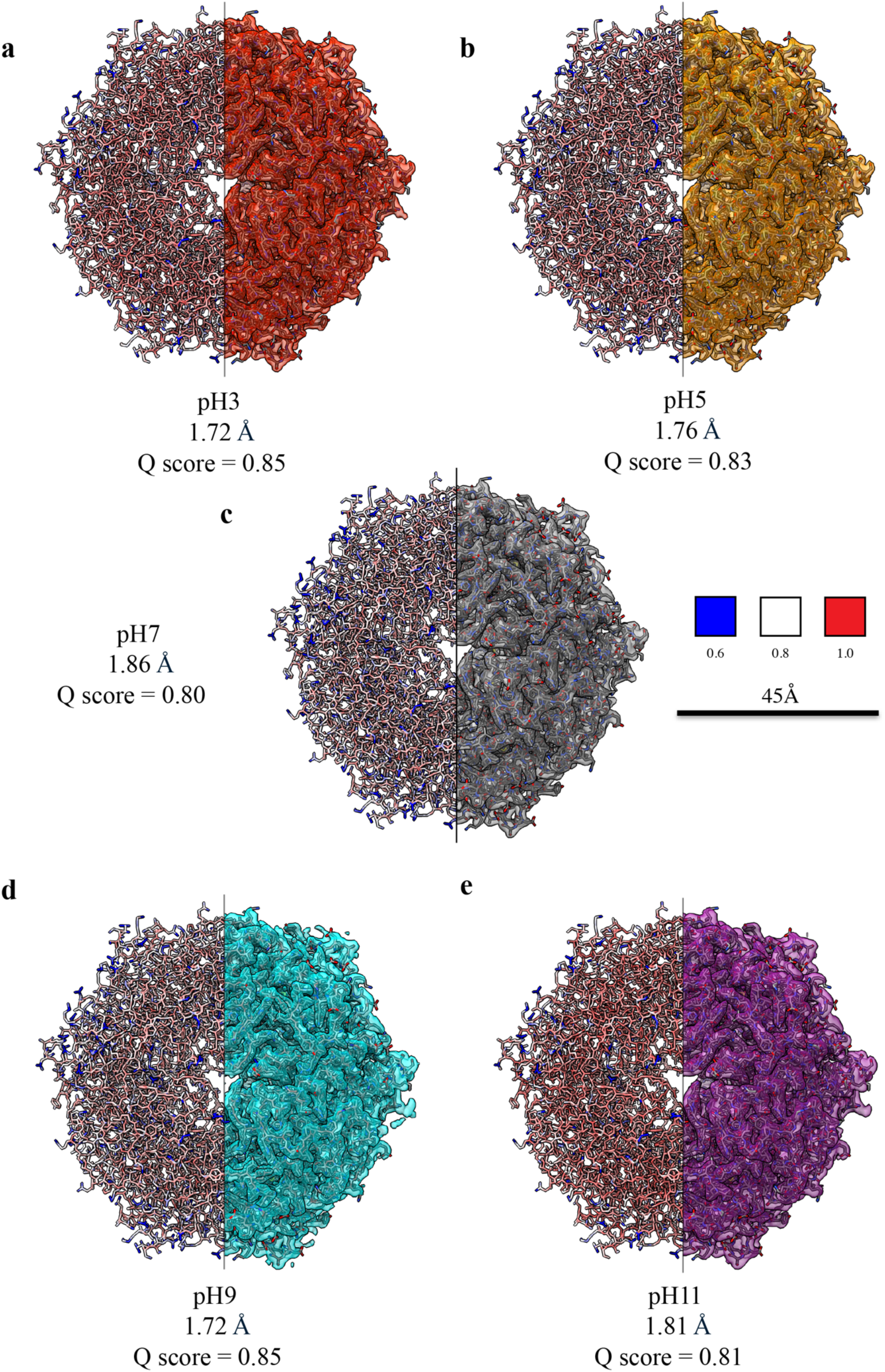
Comparison of model–map agreement across pH conditions. (**a**–**e**) Dps dodecamer structures at pH 3, 5, 7, 9, and 11, each shown with half of the particle coloured by per-residue Q-score (blue–white–red, 0.6–1.0) and the opposite half displaying the atomic model fitted into the corresponding cryo-EM density map. Reported values indicate the overall Q-score and map resolution for each reconstruction. Consistently high Q-scores (0.80–0.85) demonstrate comparable map–model quality across all pH conditions. Iron ions, present at each dimer interface, are resolved in all maps but omitted here for clarity. Scale bar 45 Å.

**Table 1.**
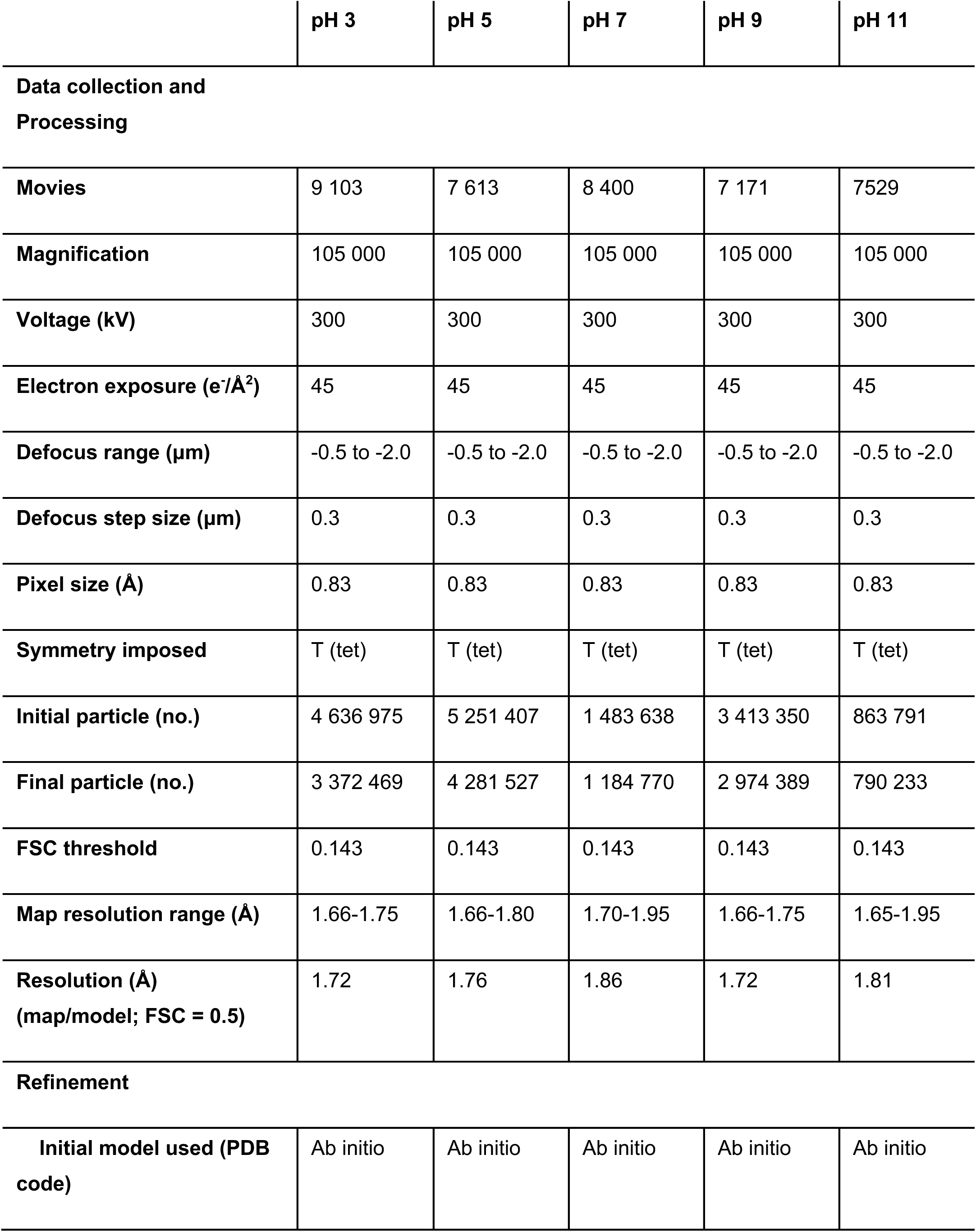
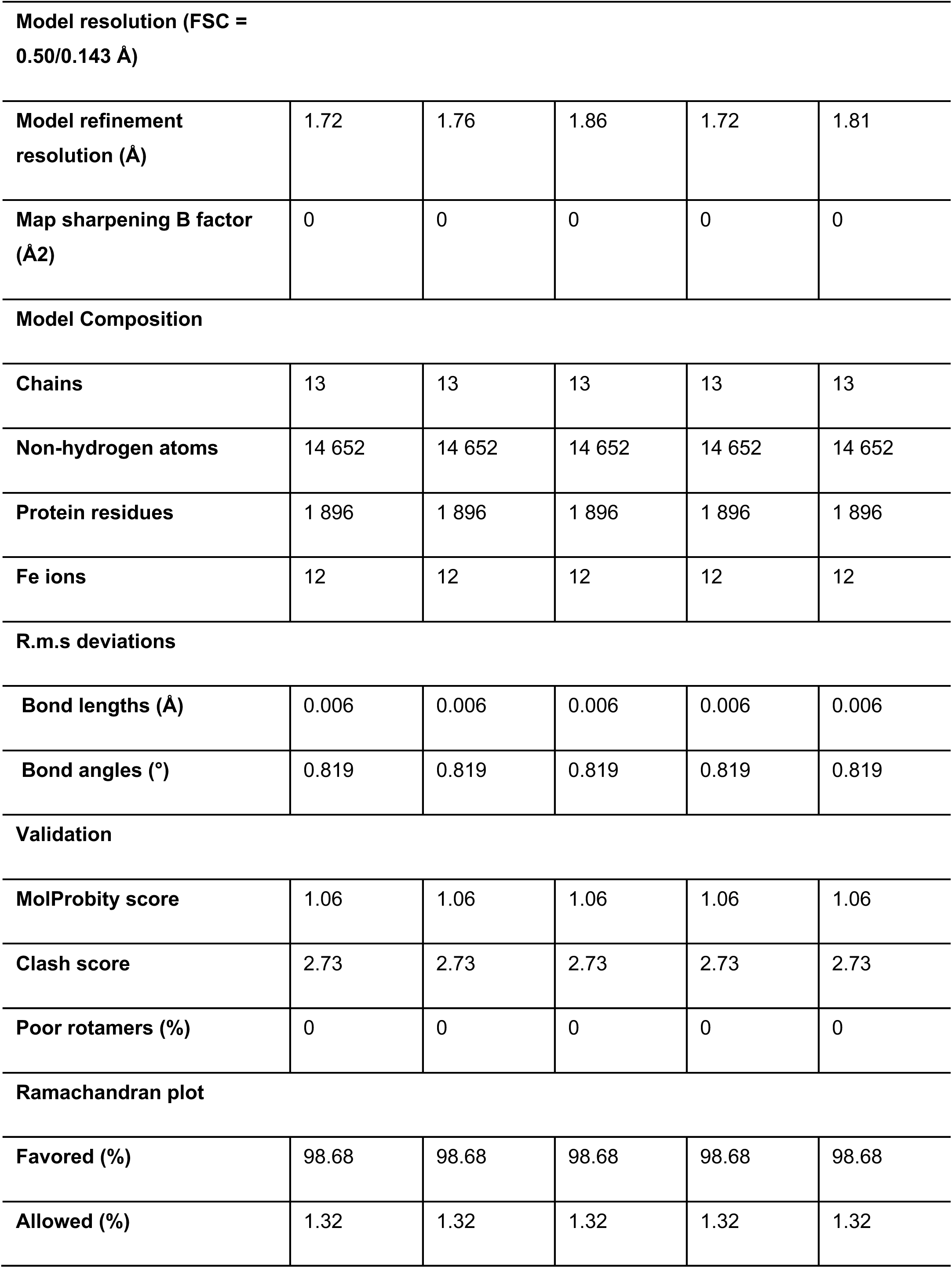
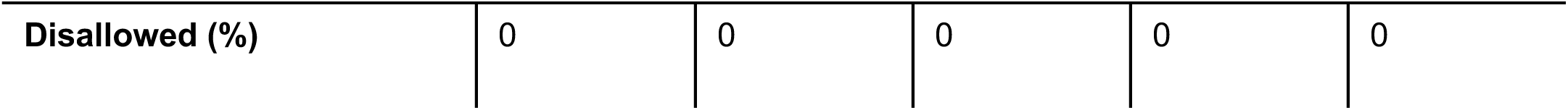
SPA and model statistics. A single refined pH 3 model was fitted into each pH-specific density map. Because the atomic coordinates were unchanged across datasets, model-based validation metrics (RMSDs, geometry, clash score, rotamers, etc.) are identical, while map-specific parameters (particle counts, FSC curves, resolutions, and sharpening factors) differ as listed. All refinement and validation statistics were obtained using ‘Phenix.real_space_refine’ and the Phenix cryo-EM validation tools [56].

|  | pH 3 | pH 5 | pH 7 | pH 9 | pH 11 |
| --- | --- | --- | --- | --- | --- |
| **Data collection and Processing** |  |  |  |  |  |
| **Movies** | 9 103 | 7 613 | 8 400 | 7 171 | 7529 |
| **Magnification** | 105 000 | 105 000 | 105 000 | 105 000 | 105 000 |
| **Voltage (kV)** | 300 | 300 | 300 | 300 | 300 |
| **Electron exposure (e<sup>-</sup>/Å<sup>2</sup>)** | 45 | 45 | 45 | 45 | 45 |
| **Defocus range (μm)** | -0.5 to -2.0 | -0.5 to -2.0 | -0.5 to -2.0 | -0.5 to -2.0 | -0.5 to -2.0 |
| **Defocus step size (μm)** | 0.3 | 0.3 | 0.3 | 0.3 | 0.3 |
| **Pixel size (Å)** | 0.83 | 0.83 | 0.83 | 0.83 | 0.83 |
| **Symmetry imposed** | T (tet) | T (tet) | T (tet) | T (tet) | T (tet) |
| **Initial particle (no.)** | 4 636 975 | 5 251 407 | 1 483 638 | 3 413 350 | 863 791 |
| **Final particle (no.)** | 3 372 469 | 4 281 527 | 1 184 770 | 2 974 389 | 790 233 |
| **FSC threshold** | 0.143 | 0.143 | 0.143 | 0.143 | 0.143 |
| **Map resolution range (Å)** | 1.66-1.75 | 1.66-1.80 | 1.70-1.95 | 1.66-1.75 | 1.65-1.95 |
| **Resolution (Å)**<br>**(map/model; FSC = 0.5)** | 1.72 | 1.76 | 1.86 | 1.72 | 1.81 |
| **Refinement** |  |  |  |  |  |
| **Initial model used (PDB code)** | Ab initio | Ab initio | Ab initio | Ab initio | Ab initio |
| **Model resolution (FSC = 0.50/0.143 Å)** |  |  |  |  |  |
| **Model refinement**<br>**resolution (Å)** | 1.72 | 1.76 | 1.86 | 1.72 | 1.81 |
| **Map sharpening B factor**<br>**(Å<sup>2</sup>)** | 0 | 0 | 0 | 0 | 0 |
| **Model Composition** |  |  |  |  |  |
| **Chains** | 13 | 13 | 13 | 13 | 13 |
| **Non-hydrogen atoms** | 14 652 | 14 652 | 14 652 | 14 652 | 14 652 |
| **Protein residues** | 1 896 | 1 896 | 1 896 | 1 896 | 1 896 |
| **Fe ions** | 12 | 12 | 12 | 12 | 12 |
| **R.m.s deviations** |  |  |  |  |  |
| **Bond lengths (Å)** | 0.006 | 0.006 | 0.006 | 0.006 | 0.006 |
| **Bond angles (°)** | 0.819 | 0.819 | 0.819 | 0.819 | 0.819 |
| **Validation** |  |  |  |  |  |
| **MolProbity score** | 1.06 | 1.06 | 1.06 | 1.06 | 1.06 |
| **Clash score** | 2.73 | 2.73 | 2.73 | 2.73 | 2.73 |
| **Poor rotamers (%)** | 0 | 0 | 0 | 0 | 0 |
| **Ramachandran plot** |  |  |  |  |  |
| **Favored (%)** | 98.68 | 98.68 | 98.68 | 98.68 | 98.68 |
| **Allowed (%)** | 1.32 | 1.32 | 1.32 | 1.32 | 1.32 |
| **Disallowed (%)** | 0 | 0 | 0 | 0 | 0 |

**Table 2.** Dps density difference statistics. Table of the statistics between the electron density maps of Dps at pH values of 3, 5, 7, 9, and 11. In each case, pH 7 was used as the reference map, with the other four compared to it using the ‘Fit to map’ function in UCSF Chimera.

| **pH** | **3** | **5** | **9** | **11** |
| --- | --- | --- | --- | --- |
| **Points** | 47 411 | 47 403 | 47 398 | 47 402 |
| **Correlation (%)** | 99.69 | 99.83 | 99.14 | 99.67 |
| **Overlap** | 339.2 | 297.4 | 337.4 | 358 |
| **Steps** | 24 | 24 | 24 | 24 |
| **Shift** | 0.00903 | 0.0026 | 0.00936 | 0.00464 |
| **Angle** | 0.00158 | 0.00251 | 0.00309 | 0.00288 |

### Structural Model and AlphaFold Predictions

An AlphaFold3 [31], prediction of the full-length Dps protein was used as the initial starting point for model building into the pH 7 cryo-EM map. Each residue along the entire length of the protein was then manually fitted into the density using COOT [32]. Any N- and C-terminal residues not visible in the density were removed. Ultimately, a validated final model comprising 12 monomeric Dps proteins spanning residues 12-169 was generated (Figure 3a). AlphaFold3 [31] predictions were then used to determine the dodecamer structure of Dps with the complete 175 AA length protein, revealing the elusive N-terminal regions as well as the C-terminal His-tagged regions that our cryo-EM maps lacked density for (Figure 3b, e). RMSD values between the experimental and AF3 data showed that the predictions were very similar in both monomeric and multimeric forms (Figure 3c, d). The 11 N-terminal residues that could not be resolved were structurally predicted to be loosely associated around the C-terminal His-tag regions, with both forming pseudo C3 symmetries, and residues at the two termini were predicted to be unordered.

**Figure 3.**
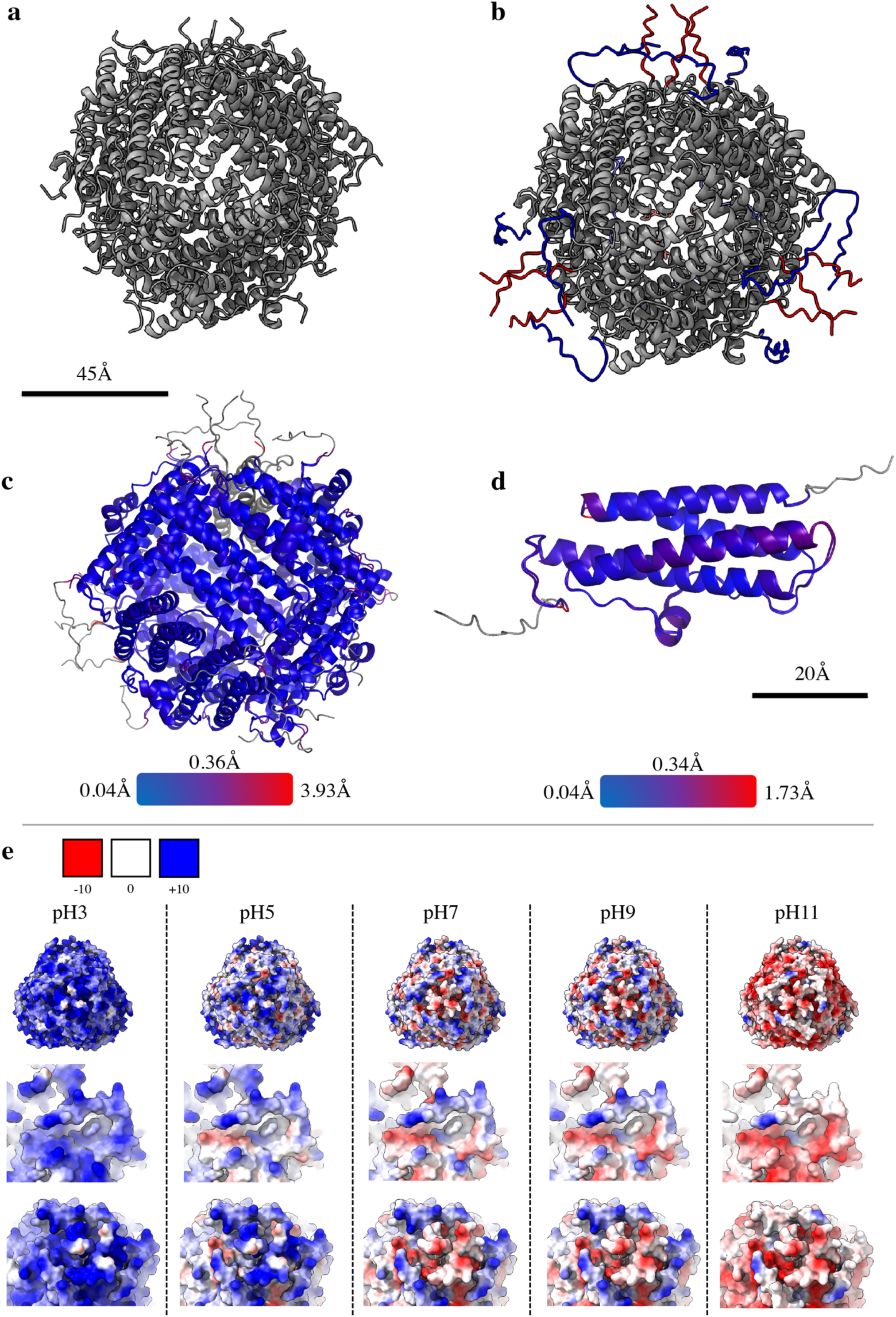
Biological/computational comparison and Electrostatic interactions. 12-mer ribbon models of Dps solved through Cryo-EM (**a**), and via AF3 [31] predictions (**b**), the latter highlighting the missing residues of the N (blue) and C (red) termini that couldn’t be solved through Cryo-EM. The RMSD values between both the solved and AF3[31] predicted Dps 12-mer (**c**) and monomer (**d**) are shown. (**e**) The electrostatic charge maps of the Dps 12-mer at pH values of 3, 5, 7, 9, and 11 (calculated via the APBS software suite [36]), where the top panel shows the AF3 predicted full-length, the middle panel highlights the N-terminus tail region, and the bottom panel visualizes the C-terminus His-tag tail region. In all instances of electrostatic charge, the range is from -10 (red) through 0 (white) to +10 (blue).

We built an atomic model of the Dps dodecamer using the pH 7 density map. Our atomic model revealed that each subunit consists of 4 α-helices. The unordered N-termini advance into the first α-helix, the largest of the 4. Upon its exit, a short, unordered bent region is found between α-1 and α-2, causing them to stack in an antiparallel manner. A very short α-helix linker region is then arranged perpendicular to α1 and joined on either side by long unordered areas, the latter of which joins to α-3 and runs parallel to α-2. A subsequent short, unordered bend region links α-3 to α-4, the latter of which is parallel in direction to α-1, before finally finishing at the C-terminal region. Both the topology of the Dps monomer and specific residue close-ups are described in Supplementary Figure 6, as well as an example of the Fe ion coordination.

Each of the twelve protomers in the Dps dodecamer contains one iron-binding site located at the two-fold (dimer) interface. The bound Fe ion is coordinated by carboxylate oxygen atoms from Asp78 and Glu78 of one subunit and by the imidazole nitrogen of His51 from the adjacent subunit, forming a mixed Fe–O/N coordination environment characteristic of Dps ferroxidase centers. Comparable architectures have been described in other Dps proteins, where conserved His, Asp, and Glu residues mediate iron binding across monomer interfaces [33, 34].

Local coordination geometry varies among the twelve sites: in all cases, two to three Fe–O interactions (Asp/Glu) are clearly resolved at 2.0–2.25 Å, whereas an Fe–N(His) contact within 2.3 Å is evident in only four of them. In the remaining eight sites, this distance is longer (≈2.3–3.2 Å) or not supported by apparent density. Consequently, no explicit LINK restraints were applied during refinement; only interactions supported by both density and reasonable distances are shown. The observed heterogeneity likely reflects a combination of subtle side-chain mobility, solvent competition, and site-to-site variation amplified by map averaging, rather than differences in Fe occupancy.

At the multimeric level, the Dps dodecamer is organized into four trimeric units, held together by C-terminal interactions between monomers. Each trimer contributes three C-terminal domains that cluster around a local threefold axis, resulting in a total of four such trimers. The N-terminal regions from neighboring monomers extend toward, and interlace around, these C-terminal trimers, such that each set of three N-termini from adjacent subunits encircles one trimer to form a pseudo-hexameric arrangement. Four of these pseudo-hexamers then associate through inter-domain contacts to produce the complete dodecameric assembly (Supplementary Figure 7). This arrangement highlights how alternating C- and N-terminal interactions cooperate to build the overall quaternary structure, with the twofold and threefold symmetry axes intertwined across the particle surface.

### The presence and absence of a His-tag in Dps

We next asked if the 6xHis-tag in our Dps construct impacted Dps-Dps interactions. Datasets at pH 5 and 7 showed that the Dps proteins formed a 2D lattice; pH 3 data also revealed frequent interactions between Dps proteins. Curious to see if the His-tag might be visible when Dps formed para-crystalline arrays, or altered the Dps-Dps local interactions, particles from the pH 3, 5, and 7 datasets were recentered onto one of the C-termini trimers as described previously in Supplementary Figure 7, before being re-extracted with larger box sizes. Rounds of 2D classification of these particles were then performed to select those interacting with neighboring Dps dodecamers (Figure 4e). Subsequent 3D classification of the particles yielded an apparent density corresponding to the C-terminal His-tag (Figure 4a). The quality of this density was insufficient for building a model capable of passing validation checks; however, a basic backbone could be traced for 2-4 of the 6 Histidine residues, depending on the monomer (Figure 4a-c). Homo refinement jobs also revealed the arrangement of Dps dodecamers relative to each other. They demonstrated that the His-tag region was oriented towards a neighboring Dps 12-mer, suggesting a physical connection between adjacent Dps multimers via the His-tags (Figure 4f). This finding was clear and in agreement with the 2D classification data (Figure 4e). The His-tag linkage corresponded to a measurement of 8 Å. The raw micrographs were also inspected to determine the X and Y distances between adjacent dodecamers, as well as the angle between them (Figure 4d).

**Figure 4.**
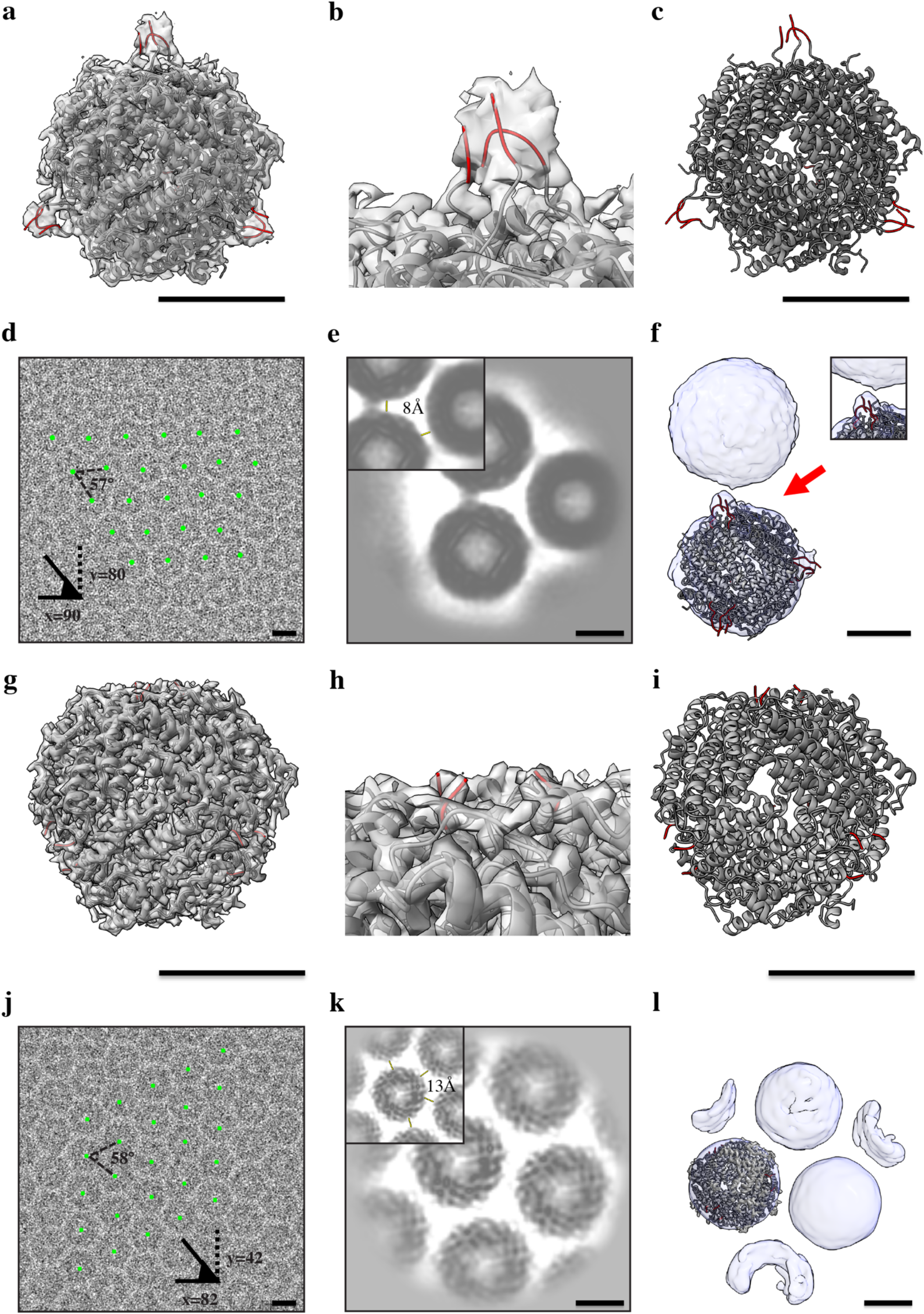
Structural comparison between tagged and untagged Dps. (**a**-**f**) data visualizing His-tagged Dps, (**g**-**l**) data visualizing untagged Dps. (**a**) volume density of the His-tagged version of Dps at pH7 with the model fitted into density, with the additionally modeled His-tag C-termini residues in red. (**b**) close-up of (**a**) to highlight the C-termini His-tag trimer (red). (**c**) model of Dps shown in **(a)** without volume density. (**d**) raw micrograph of His-tag Dps dodecamers (some centrally marked with a green dot) to visualize the tendency of His-tagged Dps at pH7 to form 2D lattice arrangements. In such lattices, each 12-mer is approximately 90 Å apart in the X-axis and 80 Å in the Y-axis from neighboring subunits, with triangles of these dodecamers separated by an angle of 57°. **(e)** 2D classification of His-tagged Dps-Dps interactions shows density between adjacent Dps 12mers and measures approximately 8 Å. (**f**) 3D volumes of particles from these 2D classes highlight the density responding to His-tag trimers, and their orientation towards a neighboring Dps 12mer. **(g)** volume density of wildtype (WT) Dps with 12mer model fitted into density, C-terminal two residues in red. (**h)** close-up of **(g)** to highlight the C-termini region and the lack of density when compared to **(b)**. **I,** model of Dps shown in **(g),** without volume density. (**j**) raw micrograph of WT Dps dodecamers (some centrally marked with a green dot) to visualize the tendency of WT Dps to form 2D lattice arrangements. In such lattices, each 12mer is approximately 82Å apart in the X-axis, 40Å in the Y-axis from neighboring subunits, with triangles of these dodecamers separated by an angle of 58°. (**k**) 2D classification of WT Dps-Dps interactions shows a lack of density between adjacent Dps 12-mers; these intermultimer distances measure approximately 13 Å. (**l**) 3D volumes of particles from these 2D classes, then highlight no density responding to any C-termini regions. Scale bar 45 Å.

The same procedures were then carried out with the data from the recently solved WT Dps structure by Sibert, Yang et al. [35]. Here, however, when the C-termini are recentered and the box size is expanded, no additional density is observed between Dps multimers in either 2D class or 3D analysis. Furthermore, the raw micrographs for this dataset yielded differing lattice X and Y values compared with those for the His-tag Dps, yet the same angle between adjacent dodecamers. These results then confirmed that the density found between the multimers of the datasets presented in this paper at pH 5 and 7 was due to His-tag residues, and that this density is responsible for the changes in inter-multimer distances of Dps. However, the angular arrangements remain the same (Figure 4g-l).

### Electrostatic Surface Shift with pH

We next investigated how the electrostatic potential of Dps changes over the pH range of 3 to 11, as this may underpin its pH-dependent DNA binding. Using the Adaptive Poisson-Boltzmann Solver (APBS) [36], we calculated the surface potential of Dps for each pH using the experimental data and AF3 predictions (Figure 3e). Electrostatic interactions were calculated for both the experimental data and AF3 predictions using the APBS suite [36], which highlighted the considerable change in surface charge Dps undergoes during the pH shift from 3 to 11. In particular, our pH predictions showed the N-terminal regions remain more positively charged than the surrounding residues across all pH values (Figure 3e). This was particularly apparent at pH 11, whereby only the N-terminal regions remain positively charged. Previous studies have shown that the N-terminal residues are responsible for DNA binding [7, 22]. Therefore, without this region of positivity, we expect DNA binding to be implausible at pH 11.

### Dps-Dps Electrostatic Interactions

To explain why Dps multimers formed a 2D lattice at pH 5 and 7, but not at the more extreme acidic pH 3 or the alkaline pH 9 and 11 (Supplementary Figure 1), we examined electrostatic interactions at the multimeric level. The data in Figure 4 confirm that lattice formations are possible both with and without His-tagged proteins; however, the details of these protein-protein interactions remain elusive. Our electrostatic computational data showed that the Dps dodecamers at pH 3 have a uniform, very positive charge (Figure 3e). The concentrations of Dps samples frozen at each pH value were identical, as were the freezing conditions; thus, we concluded that, as the only changed parameter, the electrostatics due to pH shifting is the cause of 2D lattice formation. The electrostatic data at pH 11 were consistent with this hypothesis, showing sufficiently negative values, and were supported by micrograph observations, which indicated that pH 11 Dps was also incapable of forming 2D crystal structures (Figure 3e).

Electrostatics of Dps at pH 5, 7, and 9 show favorable interactions, with sections of the Dps dodecamer appearing positively and negatively charged, suggesting that Dps-Dps attractions are feasible. Indeed, these computational calculations were supported by experimental data at pH 5 and 7 (Supplementary Figure 1). A close inspection of the pH 9 dataset revealed subtle quantities of non-uniform material between Dps multimers. It was unclear whether this content was a denatured Dps protein or a result of salt interference at the high pH. In any case, we suspect that, despite electrostatics theoretically allowing for Dps-Dps lattice interactions, potential salt aggregates or protein denaturation can alter these lattice-forming interactions. Similar denaturation/salt artifacts are also seen at pH 11 (Supplementary Figure 1); however, at pH 11, the electrostatic data would suggest that Dps-Dps binding is unfavorable, and so even if denaturation at this pH value were not present, we still wouldn’t expect to witness lattice formation within these micrographs.

### Dps Binding Affinity is Maximal at pH 5 and Drops at pH 11

As mentioned, EHEC is subject to pH shifting orchestrated by the stomach acid during digestion. As intracellular matrix pH decreases in *E. coli* cells, Dps is overexpressed in response and subsequently binds DNA in large quantities to protect it from denaturation [6, 7, 37]. With this in mind, we sought to determine the pH range over which Dps-DNA binding might operate. To this end, we conducted DNA-binding assays at pH 3, 5, 7, 9, and 11. Fixed amounts of DNA were mixed with Dps at pH 11, with concentrations ranging from 0 to 1000 ng and from 0 to 3500 ng. These mixtures were run on 0.5% agarose gels in triplicate and subsequently imaged for analysis (Figure 5b, Supplementary Figure 8). Densitometry-derived fraction bound values were fit with a 4-parameter logistic model, yielding 𝐸𝐶_50_ values (the concentration at half of the fitted Top). Because the Hill slope deviates from 1 for some conditions, we report 𝐸𝐶_50_ (K½, apparent) rather than an authentic K_d_. Our results indicate that the binding affinity of Dps-DNA is strongest at pH 5, with an 𝐸𝐶_50_ value of 73.4 nM (Figure 5a, Table 3). Successful activity was also observed at pH levels of 3, 7, and 9, with 𝐸𝐶_50_ values of 185.5 nM, 165.4 nM, and 224.9 nM, respectively. Indeed, Dps-DNA interactions are still observed even at pH 11; however, with an 𝐸𝐶_50_ value of 815.5 nM, the ability of Dps to bind DNA at pH 11 is approximately 8 times lower.

**Figure 5.**
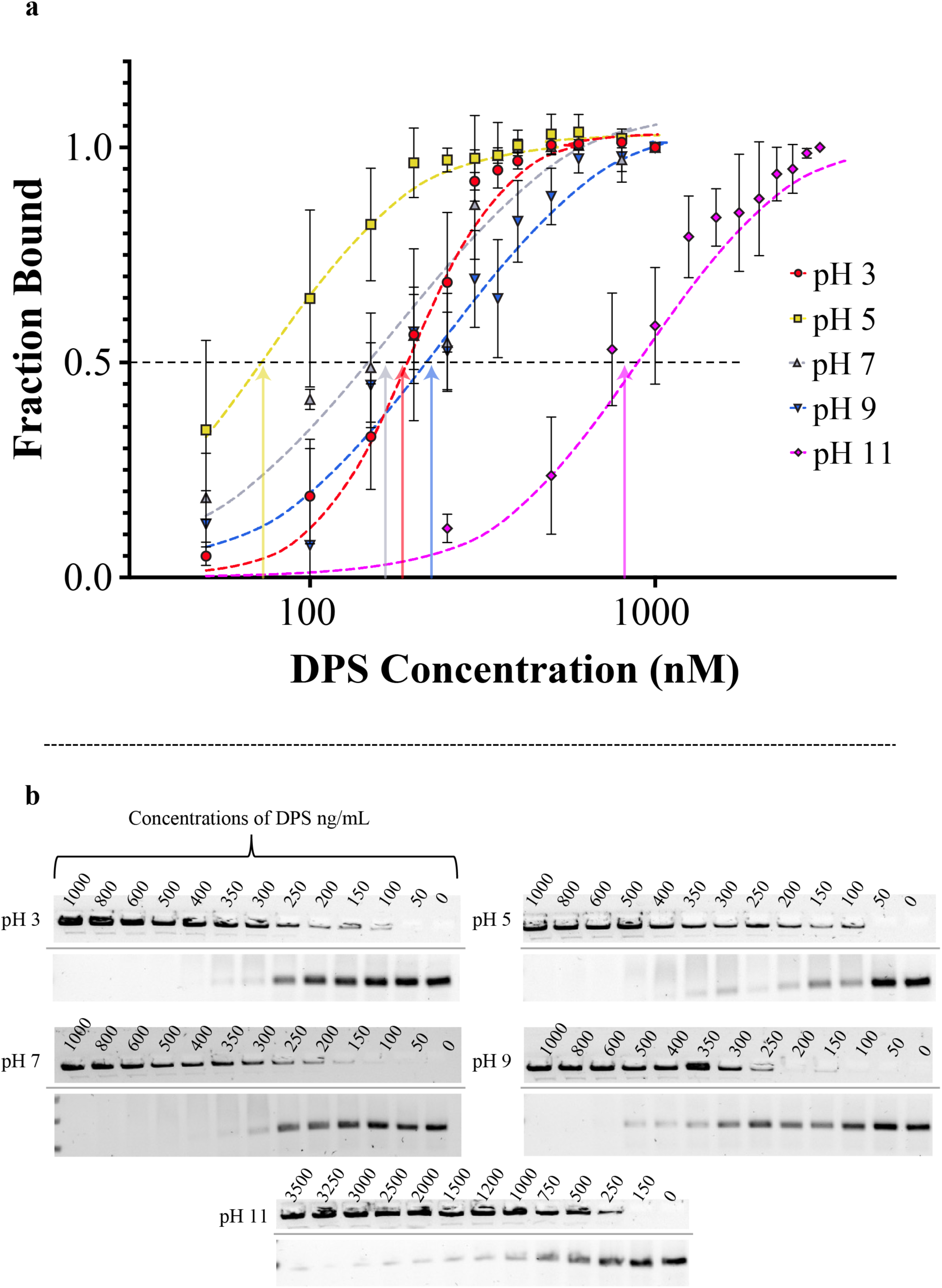
pH-dependent DNA binding and compaction by Dps. (**a**) Quantification of Dps–DNA binding from electrophoretic mobility shift assays (EMSAs). Each curve shows the mean fraction of DNA bound () ± SD) across three independently prepared and imaged gels that passed predefined quality control criteria (monotonic trend, near-saturation at high concentrations, and uniform background). For each pH, the value was calculated from the loss in free-DNA band intensity, normalized to the near-saturating 1000 nM Dps lane. Curves represent nonlinear regression fits to a four-parameter variable-slope Hill model (GraphPad Prism 10, ‘*[Agonist] vs Response — Variable Slope’*). The *Bottom* plateau was constrained to 0, while *Top*, 𝐸𝐶_50_, and the Hill coefficient (𝑛) were free parameters. The dashed line denotes 50 % bound (apparent 𝐸𝐶_50_), and each of the colored arrows shows the 𝐸𝐶_50_ values for the relevant pH value: 𝐸𝐶_50_ pH 3 = 185.5 nM, pH 5 = 73.4 nM, pH 7 = 165.4 nM, pH 9 = 224.9 nM, and pH 11 = 815.5 nM. (**b**) Representative agarose DNA gels showing Dps-dependent DNA binding at the indicated pH values, whereby only the tops and bottoms of each gel are shown (deleted region marked by grey lines). Each lane contained identical quantities of 448bp DNA, with increasing Dps concentrations (0–1000 nM for pH 3,5,7,9, and 0-3500nm for pH 11 above each lane). Dps binds DNA and halts its migration, resulting in reduced free-DNA band intensity near the bottom of the gel and, at high concentrations, complete retention of DNA in the wells. The degree of retention increases with decreasing pH, consistent with enhanced electrostatic interactions between Dps and DNA under acidic conditions.

**Table 3.**
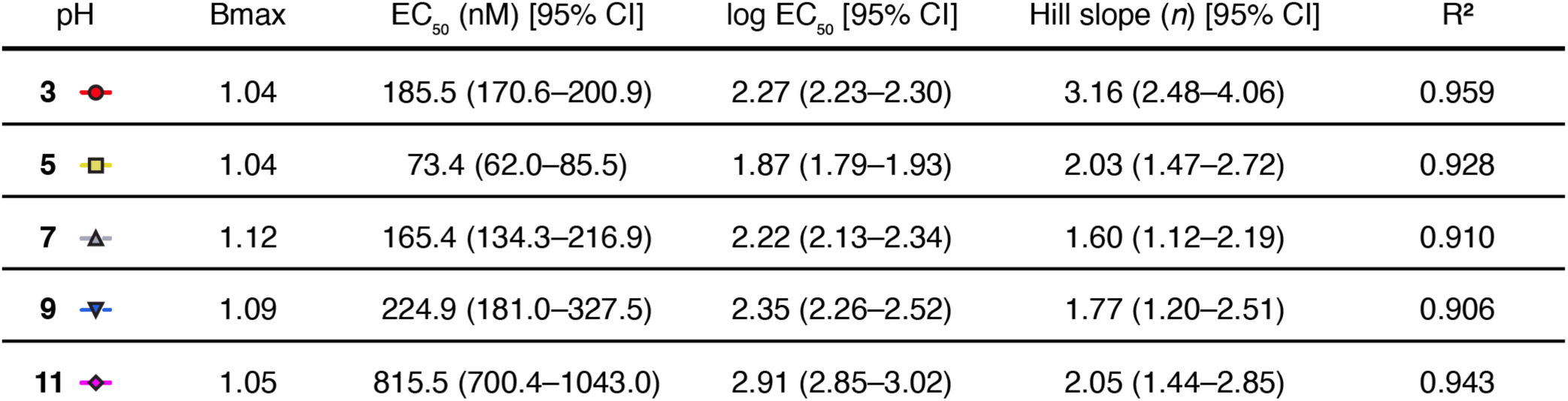
Summary of Hill-model parameters describing Dps–DNA binding across pH 3–11. Best-fit parameters obtained from nonlinear regression of DNA binding data (Figure 5) to a four-parameter variable-slope Hill equation in GraphPad Prism 10 (*[Agonist] vs Response — Variable Slope*). Each dataset (pH 3–11) was fit independently using averaged fraction-bound values from three replicate gels (mean ± SD). The *Bottom* plateau was constrained to 0, while *Top* (𝐵_max_), 𝐸𝐶_50_, and the Hill slope (𝑛) were free parameters. Values represent the best fits, with 95% confidence intervals (CIs) derived from profile likelihood estimation. 𝐸𝐶_50_ values report the apparent Dps concentration required for half-maximal DNA binding. The lower values (pH 3–5) indicate a stronger binding affinity under acidic conditions, while binding progressively weakens at higher pH (≥ 9). All fits showed high goodness-of-fit (𝑅^3^ ≥ 0.90).

| pH | Bmax | EC<sub>50</sub> (nM) [95% CI] | log EC<sub>50</sub> [95% CI] | Hill slope (*n*) [95% CI] | R<sup>2</sup> |
| --- | --- | --- | --- | --- | --- |
| 3 | 1.04 | 185.5 (170.6–200.9) | 2.27 (2.23–2.30) | 3.16 (2.48–4.06) | 0.959 |
| 5 | 1.04 | 73.4 (62.0–85.5) | 1.87 (1.79–1.93) | 2.03 (1.47–2.72) | 0.928 |
| 7 | 1.12 | 165.4 (134.3–216.9) | 2.22 (2.13–2.34) | 1.60 (1.12–2.19) | 0.910 |
| 9 | 1.09 | 224.9 (181.0–327.5) | 2.35 (2.26–2.52) | 1.77 (1.20–2.51) | 0.906 |
| 11 | 1.05 | 815.5 (700.4–1043.0) | 2.91 (2.85–3.02) | 2.05 (1.44–2.85) | 0.943 |

### Cryo-ET Reveals Globular Dps–DNA Assemblies on Short DNA

The segmentation data shown in Figure 6 illustrated the straightforward 3D amalgamation of Dps at pH 7, which appeared structurally distinct from the 2D lattice formations seen in the absence of DNA (Supplementary Figure 9c). Previous cryo-ET studies concluded that Dps forms crystalline 3D lattice structures around DNA from *E. coli* [9, 17]; however, those experiments used DNA lengths of 9,900 bp. In contrast, the DNA used in our Dps-DNA experiments was 448 bp and comprised non-specific, non-coding regions not derived from *E. coli* (Supplementary Figure 10b, c). This design choice aimed to test whether *E. coli* Dps displays sequence or length preferences. Prior publications have been conflicted as to whether Dps exhibits specificity for certain DNA regions, shapes, or conformations [7, 26, 38]. To further explore this, we incubated *E. coli* Dps with 448 bp DNA fragments from *Rhodobacter sphaeroides,* which were previously generated in prior studies [39]. Both the tomographic globoid structural data and the Dps-DNA binding curve data (Figures 5, 6, Supplementary Figure 8, 9) confirmed that *E. coli* Dps binds non-specifically and can associate with foreign genomes.

**Figure 6.**
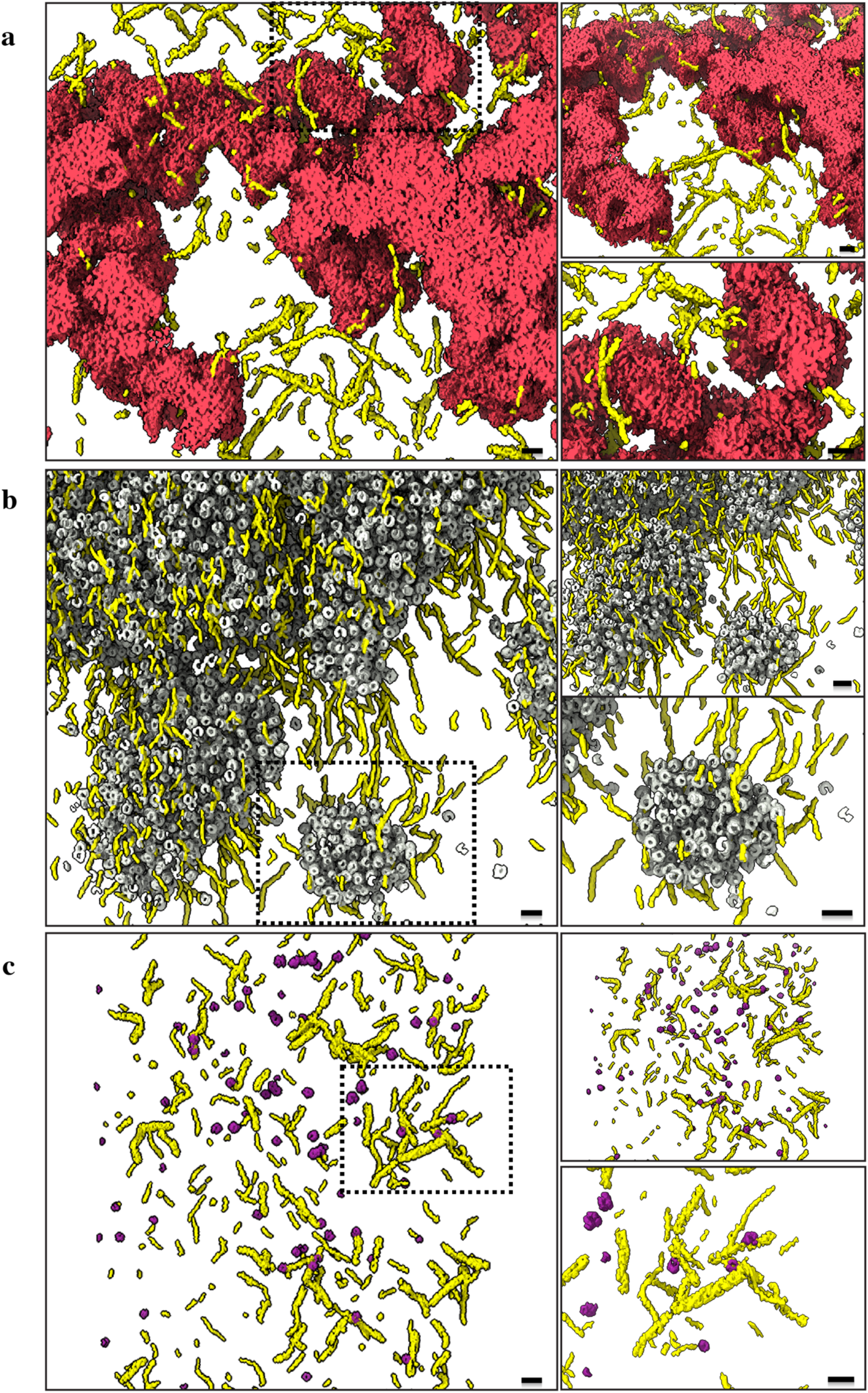
Dps-DNA segmentation. (**a**) Segmentation of Dps-DNA interactions at pH 3, where Dps are shown in red and DNA in yellow. A globular mass of Dps can be seen with DNA filaments sticking out from a front view (left), as well as a 30° rotation about the X axis to illustrate the depth of the Dps-DNA globoids (top right). A zoomed view shows the non-specific Dps-DNA interactions in the bottom-right corner. (**b**) Segmentation of Dps-DNA interactions at pH7, where DPS are shown in grey and DNA in yellow. Dense collections of Dps individual multimers can be seen sequestering around DNA from a front view (left) and a 30° rotation about the X axis, illustrating the depth of the Dps-DNA globoids (top right). A zoomed view of the smaller Dps-DNA globoid, highlighted in the dotted box, is shown in the bottom-right panel, underscoring the DNA strands protruding from the Dps assemblage. (**c**) Segmentation of Dps-DNA interactions at pH11, where Dps are shown in purple and DNA in yellow. Sparse collections of Dps individual multimers can be observed within a population of DNA strands, as seen from a front view (left) and a 30° rotation about the X axis (top right), to illustrate the lack of interaction between Dps and DNA. A zoomed view of the smaller Dps-DNA local interactions highlighted in the dotted box can be seen in the bottom right, showing the DNA strands and how they aren’t heavily engaged with Dps monomers, and thus don’t form globoids. Scale bars are 200 Å.

To examine how pH influences Dps-DNA interactions, cryo-ET was performed on samples incubated at pH 3, 7, and 11, conditions chosen to represent the maximal, median, and minimal pH ranges, respectively, as informed by our assay data (Figure 5). After incubation, Dps-DNA mixtures at each pH were plunge-frozen for tomographic data collection suitable for sub-tomogram averaging (STA) and segmentation. Because fiducial markers tended to aggregate at extreme pH values, samples were generated without addition of any fiducial particles, and the best tilt series from each condition were reconstructed using AreTomo [40], to generate bin6 tomograms. These image stacks were then processed through the IsoNet [41] pipeline to enhance contrast and partially correct for missing-wedge artifacts. Videos were generated from these tomograms; supplementary videos 1, 2, and 3 correspond to the tomography data at pH 3, 7, and 11, respectively. Subsequent segmentation was then performed using DragonFly [42], with supplementary videos 4, 5, and 6 created, in which independent models were trained on the pH 3, 7, and 11 datasets to avoid segmentation bias.

Segmentation of DNA was markedly less accurate than that of Dps, as DNA is a flexible, structurally unordered strand that can adopt numerous conformations. Dps, by contrast, are large, spherical, and uniform in morphology, making them easily identifiable through Z-slices of the tomograms. Consequently, manual picking and model training within DragonFly could reliably segment Dps but not most DNA; only DNA strands extending from or adjacent to Dps clusters were confidently identified. Nevertheless, the raw tomograms clearly revealed DNA filaments traversing through Dps clusters, particularly at pH 7 (Supplementary Figure 9d).

At pH 7, 12 tilt series of Dps-DNA interactions were recorded, yielding eight high-quality tomograms in which both Dps and DNA are readily visible. In these reconstructions, Dps 12-mers formed large globular assemblies that encapsulate DNA strands (Figure 6b, Supplementary Figure 9d). Simple measurements of these tomograms revealed that the Dps-DNA interactions formed two major phenotypes: large and small globoids. The large globoid structures displayed thousands of Dps 12-mers interacting to form an entity with no obvious breaks, often measuring over 200 nm in the X and Y axes and approximately 120 nm in thickness along the Z axis. Indeed, many DNA strands were seen poking out of these vast structures, and cross-sectionally within the spaces between Dps.

Conversely, the small globoid structures mimicked this same design, but instead were found isolated away from the large globoids. These small globular structures were often measured to be approximately 100 nm in the X and Y axes and 75 nm in the Z axis. Figure 6b and Supplementary videos 2 and 5 visualise both phenomena within a single tomogram and segmentation. Within Figure 6b, a dotted black box denotes the small globoid, and it is distinctly separate from the large globoid above it. This segmentation and tomography data clearly emphasize that Dps at neutral pH organize into globular complexes rather than the crystalline lattices previously reported.

Comparatively, the pH 3 data exhibited lower clarity; although both DNA and Dps were identifiable (Supplementary Figure 9b), they were less distinct. Given these limitations, segmentation focused on visualizing entire Dps structures rather than individual Dps 12-mers (Figure 6a). This approach revealed Dps-DNA different in size and morphology from those seen at pH 7. Instead of consistent globular shapes composed of Dps, with DNA strands weaving throughout the gaps between them, at pH 3, we observed Dps forming thinner and longer interactions reminiscent of tubular-type structures. The approximate diameter of these structures was 40-50 nm and extended in length across both the Y and Z axes to form complex shapes. This ultimately created Dps-DNA interactions with holes and craters that weren’t seen at pH 7, and instead of globoid shapes, more tubular designs were found (Supplementary video 1 and 4). Approximately 52 tilt series were collected across two grids to assess ice quality, and 8 high-quality tomograms were further processed.

At pH 11, 12 tilt series were acquired, yielding six well-resolved tomograms. Similar to pH 7 datasets, both Dps 12-mers and DNA strands were clearly observed (Figure 6c, Supplementary Figure 9f); however, minimal interaction between the two populations was evident, and no globular assemblies were detected. Visually, this resulted in both DNA strands and Dps 12-mers being observed randomly dispersed within the tomograms (Supplementary video 3 and 6).

Collectively, these results demonstrate that Dps readily formed large assemblies with short, non-specific DNA at pH 3 and 7, but failed to do so under alkaline conditions. The observed correlation between these structural states and the binding-assay data underscores the pH-dependent modulation of Dps-DNA interactions and the structural adaptability of Dps in mediating genome protection across diverse environmental conditions (Figures 5 and 6, Supplementary Figures 8 and 9).

## Discussion

### Dps Maintains Structure But Changes Interactions Across pH

Our comprehensive analysis reveals that *E. coli* Dps remains structurally unperturbed in its tertiary and quaternary structure across a pH range of 3 to 11. This extreme stability is consistent with Dps’s role as a long-term protector of DNA during dormancy or stress, further highlighting that Dps must remain intact even when cytosolic pH fluctuates or the environment becomes acidic. These findings are in line with earlier biophysical studies on *M. smegmatis* Dps, which reported reversible dimerization at pH values below ∼6 and above ∼7.5 [43, 44]. Interestingly, while our high-resolution imaging data suggest the dodecamer persists, the micrographs and 2D class averages gave hints of pH-dependent higher-order behavior: at pH 7, we saw Dps forming crystalline-like lattices on the grid (Figure 4c, Supplementary Figure 1) similar to Antipov *et al.*’s observations [27], but at pH 3, those lattices were completely absent, replaced by irregular clusters of Dps. At pH 11, many Dps particles appeared separate, and some showed signs of partial unfolding with elongated or smeared particles in the micrographs or in the 2D classes. We interpret this as evidence that Dps–Dps interactions are also pH-sensitive. A neutral pH favors a weak, attractive interaction between Dps shells, which may be mediated by complementary charge patches, allowing them to align in 2D or 3D arrays when concentrated. A low pH (pH 3) likely causes nonspecific aggregation, in which Dps shells adhere to one another in a disordered manner, possibly due to the overall positive charge and reduced electrostatic repulsion. High pH (pH 9–11) renders the protein shells net negative, introducing strong repulsion that prevents both ordered and disordered aggregation, while simultaneously destabilizing the subunit interfaces, releasing some monomers. Altogether, the apparent crystallization of Dps is a pH-dependent phenomenon, driven by electrostatic charge states.

The current assumption within the literature is that Dps categorically form crystal structures around DNA [5, 9, 22, 25, 28]. However, all evidence has been conducted at pH 7 only. At least for EHEC strains, the ability of these pathogenic cells to survive at pH 1 is plausible, given the high concentrations of stomach acid in humans. Whilst the cytoplasmic pH of EHEC under these conditions would never get down to a value of 1, their internal pH would certainly become acidic. Prior cytosolic pH probe experiments with *E. coli* have demonstrated that an external pH of 3.0 yields a cytosolic pH of 4.4 [45]. As observed in SPA cryo-EM micrographs, Dps 2D crystallization is patchy at pH 5, and it’s not possible at pH 3. This then begs the question: are the published 3D Dps-DNA crystal events capable of being replicated under the acidic conditions that EHEC faces? Or is the crystallization not paramount to EHEC’s survival?

### Local vs. Global DNA Organization – A Two-Stage Model

A central finding of this work is that short DNA segments of hundreds of base pairs, bound by Dps, do not form lattice structures at any pH tested. Instead, we saw local “globoids” or tubular arrangements of Dps dodecamers with DNA weaving through at pH 3 and 7 (Figure 6a, b). This suggests that Dps can compact DNA locally without creating a long-range order. Only when DNA exceeds a specific length or is densely packed can local clusters merge into a crystal lattice. Prior work also supports this view; Wolf *et al.* first postulated that Dps–DNA co-crystals form on chromosomal-length DNA [25]. Later, Dadinova *et al.* demonstrated *in vitro* co-crystallization using ∼10 kbp DNA, but were unable to achieve it with short oligonucleotides [9]. Our data support the notion that DNA length or topology is the primary determinant of whether Dps assembles into a periodic lattice or an amorphous aggregate.

We propose a two-stage assembly model for Dps in the nucleoid. Stage 1: Nucleation, where Dps dodecamers bind along DNA, neutralizing charge and compacting segments into tiny clusters, which we observe as globoids by cryo-ET (Figure 6). This stage is rapid, driven by electrostatics, and could be accelerated by a lower pH and higher Dps concentrations. Stage 2: Ordering, if conditions allow, such as a physiological pH ∼7, relatively low ionic strength, and crucially, long contiguous DNA, the Dps–DNA clusters may rearrange into a crystalline array. DNA acts as a scaffold or template in this ordering, as evidenced by straight DNA filaments seen between Dps particles in crystals. In this manner, DNA bridges Dps units at specific intervals, promoting a regular spacing. Our results show that in the absence of sufficient DNA length, stage 2 doesn’t occur, and the system remains in an amorphous and more “phase-separated” state. This has biological implications, as the cellular environment may support the organization of the nucleoid into loops and domains. Dps might fully crystallize in regions where multiple DNA loops come together and Dps is highly concentrated, while other areas remain semicondensed, non-crystalline. This heterogeneity was indeed observed by Loiko *et al.* in *E. coli* under starvation conditions [46].

To explain the difference in Dps-DNA morphologies observed in the tomography data, we must consider the ionic interactions at play. At pH 7, we observe large, well-defined globoids of Dps entwined with DNA, consisting of tightly packed, roughly spherical condensates that correspond more closely to the nanocrystalline state documented in starved bacteria [46]. The apparent difference is that short DNA prevents proper crystallization from occurring. At pH 3, however, Dps still cluster with DNA, but the morphology shifts dramatically; instead of full globoids, we see irregular, tubular, or blob-like networks with distinct holes and less densely packed Dps. We propose that acidification alters the balance of Dps–Dps and Dps–DNA interactions. Such actions do not prevent binding, but disfavor the final self-lock-in stage necessary to form the compact crystalline globules [22]. In this scenario, Dps–DNA assemblies default to a more open ‘folded nucleosome-like’ architecture—structures previously observed when Dps packing is impaired [46]. Thus, the pH 3 condensates we imaged likely reflect functional engagement but structurally sub-optimal assembly: sufficient to bind DNA, but unable to collapse into the dense globoids seen at physiological pH values.

### Effect of pH on *In Vivo* Dps Function

Another motivation for this study was to simulate the intracellular pH range that pathogens or soil bacteria might experience and monitor Dps function. *E. coli* O157:H7, for example, can survive stomach transit by acid tolerance mechanisms [37]. In such cases, the pH of the periplasm and cytoplasm can drop to approximately pH 4.5 [45, 47]. In line with our findings, at pH 4.5, Dps is likely to be predominantly in its protonated, DNA-bound form, consistent with our observations at pH 5. We demonstrated that at pH 5, Dps binds DNA strongly (Figures 5 and 6, Supplementary Figures 8 and 9) and, in the absence of DNA, forms two-dimensional paracrystalline arrays (Supplementary Figure 1). By pH 3, crystalline order was absent; Dps was dispersed and presumably adhered to DNA or other Dps dodecamers in a disordered manner. This raises an interesting question: Does Dps need to form an ordered crystal on DNA to protect it, or is a disordered condensate sufficient? In the context of acid stress, as seen in EHEC, it is possible that a full biocrystal is not formed when the cytosolic pH may be too low for long-range order. Yet, Dps still protects DNA by coating and coalescing with it. Indeed, the study by Jeong *et al.* found that, *in vitro*, Dps protected DNA at pH levels of 2.5–3 [37], presumably without forming a crystalline structure. Therefore, crystallization *per se* may not be paramount to survival under acidic conditions; it could be a side effect observed mainly during starvation at neutral pH. This is interesting when considering the role of Dps, where the assumption has been that Dps always forms crystals around DNA at the stationary phase of bacterial growth [25]; however, we suggest that this might be conditional. Under acidic conditions, Dps–DNA complexes might remain amorphous yet still confer protection. Testing Dps–DNA assembly at different pH in cells, perhaps via correlative light and electron microscopy (CLEM) imaging or crosslinking approaches, would be very illuminating.

### Electrostatic Driving Force

Our data support the classic view that electrostatics are the primary driver of Dps–DNA association [29]. The N-termini remain positively charged over a wide pH range and are undoubtedly responsible for the initial attraction to DNA [48]. The correspondence between our APBS-calculated surface potential (Figure 3e) and the EMSA affinities is striking (Figure 5a and Table 3). At a low pH of 5, the entire protein is positively charged, which correlates with strong binding (*EC*_50_ ∼73 nM). At a high pH of 11, the protein is predominantly negatively charged, except for the faintly positive tails, which correlate with weaker binding (*EC*_50_ ∼815 nM). Furthermore, the cryo-EM structures reveal no conformational change that could account for the binding difference, indicating that Dps does not exhibit open or closed states; rather, the charge state is the determining factor. Thus, Dps does not appear to have an allosteric pH switch. Instead, it seems to function as a quasi-static electrostatic machine, binding DNA when oppositely charged and releasing it when like-charged.

This led us to question whether there are any secondary contributors to Dps binding with DNA. Hydrophobic interactions or shape complementarity might play minor roles. For example, the N-terminus of Dps contains several hydrophobic residues (e.g., Met1 and Ala4 in *E. coli*) that could transiently interact with the minor groove of DNA or its sugar backbone. There is also the potential role of bridging multivalency, as each Dps has 12 ‘arms’ that allow cooperative binding. In our binding curves, the Hill coefficients ranged from 1.60 to 3.16. Thus, this strongly suggests a positive correlation, indicating that initial binding of Dps-DNA promotes further Dps-DNA interactions in a localized manner. This cooperativity effectively sharpens the binding transition, such that once a threshold is reached (e.g., sufficient protonation, sufficient Dps concentration), Dps rapidly saturates the DNA. This behavior is consistent with the formation of the condensed complexes we observed (Figure 5).

### His-Tagged Dps vs. Wild-Type

Another outcome of our study is the validation that a C-terminal His-tagged Dps retains wild-type behavior across all assays. Structurally, our cryo-EM maps of His-tagged Dps at pH 7 matched the 1.6 Å crystal structure of untagged Dps (PDB 1DPS) with minimal differences [5]. Functionally, we observed that His-tagged Dps binds to DNA and forms 2D and 3D lattices, consistent with reports for native Dps [49]. This is notable because a previous report [43] found that attaching a His-tag to the C-terminal tail of *M. smegmatis* Dps, which binds DNA, significantly enhanced DNA condensation at neutral pH. Essentially, the positively charged His-tag was possibly an extra “glue” and caused the protein to condense DNA into clusters more readily. In our case, the C-terminus of *E. coli* Dps is not known to participate in DNA binding; instead, the functional end is the N-terminus. Our EMSA results did not indicate any aberrant higher-order aggregation at neutral pH; in fact, they show a gradual binding curve (Figure 5a). We also confirmed that Dps with the His-tag formed crystals, as noted in the 2D Dps lattice at pH 7 (Figure 4d, Supplementary Figure 1), which was with the His-tagged protein. Therefore, a key takeaway for researchers is that a C-terminal His-tag on *E. coli* Dps is potentially benign in terms of its structure and DNA-binding function. The incorporation of the His-tag facilitates future work and could even be potentially valuable for biotechnological applications of Dps.

### Physiological Implications and Future Directions

Our results highlight that Dps acts as a pH-responsive nucleoid compactor. In an *E. coli* cell entering the stationary phase, the cytosol may become slightly acidic, with a change from a pH of 7.6 during logarithmic growth to ∼7.0 in the stationary phase, or lower under conditions of nutrient deprivation. This change would increase the DNA affinity and promote nucleoid binding of Dps. Simultaneously, competitors like IHF, which prefer neutral or alkaline conditions, would bind less avidly [29]. Thus, environmental cues feed into nucleoid organization through simple physicochemical rules. The fact that Dps still binds DNA at neutral pH also ensures that once it is more abundant in the cell, it will inevitably bind and occupy DNA, even if the overall pH hasn’t dropped [11].

An intriguing application to consider is the design of pH-sensitive DNA protection systems. Dps or engineered variants could be used to package DNA or plasmids for delivery, with release triggered by pH shifts, such as an acid-triggered DNA release, which might be helpful for cargo delivery in acidic organelles. Conversely, one could harness Dps to protect DNA in acidic environments, which could already be relevant for probiotics or oral DNA vaccines that endure stomach acid.

Within the context of the cell, it would be interesting to explore how Dps-mediated DNA compaction interacts with other cellular processes. Do processes associated with Dps DNA condensation entirely halt transcription? This is plausible given that Dps represses transcription globally, as shown by its global repression effect [7]. If this is the case, cells may already be programmed to control the onset of Dps binding, perhaps by modulating the transient nature of Dps-DNA interactions under specific pH or ionic conditions. For example, suppose a stationary-phase cell encounters a slight improvement in conditions with rising pH and/or nutrient levels. In that case, Dps might partially disengage, allowing gene expression to proceed, as seen in the so-called “Dps-null” effect, where gene expression patterns change in the absence of Dps [7]. The competitive binding between Dps and other NAPs, such as IHF or HU, is also an area ripe for biophysical study. Our work provides a baseline for Dps alone, and combining Dps with IHF on DNA at various pH levels could reveal how they dynamically partition the genome [29].

In conclusion, by examining Dps at the atomic and mesoscale assembly levels under various pH conditions, we have provided a unified view of how this protein functions as an adaptable DNA-packaging machine. Dps’s ability to maintain its structure while tuning its DNA binding with pH underscores its elegance as an evolutionary solution for protecting the genome when life becomes hostile.

## Methods

### DPS plasmid construction

Plasmid pKCJ0325, which carries the Dps gene, was kindly donated to us by the Kaspar group [37]. From here, amplification was performed to generate pET21b-DPS, which was ultimately used for Dps expression. The resulting plasmid was then verified by restriction digest analysis and whole plasmid sequencing using Plasmidsaurus (Plasmidsaurus, Eugene, OR) (Supplementary Figure 10a).

### DPS protein expression and purification

For protein expression, pET21b-DPS was transformed into *E. coli* BL21(DE3). Induction of protein expression was carried out by inoculating 500 mL of LB broth, supplemented with ampicillin at 100 mg/mL, with 2.5 mL of an overnight culture of BL21(DE3) harboring pET21b-DPS. This culture was grown to an optical density at 600 nm (OD_600_) of ∼0.5 at 37°C and 200 RPM shaking, and IPTG-induced Dps expression was initiated by adding 1 mM IPTG. DPS expression was induced for 4 hours, then the cells were pelleted at 5,000 × g for 10 minutes at 4°C. The supernatant was discarded, and the cell pellet was stored at -80°C until purification.

Dps purification was performed as follows. The cell pellet was resuspended in 10 mL of buffer: 10 mM Tris at pH 6.0, 100 mM NaCl, 100 μL of 20 mg/mL lysozyme, and 200 μL of protease inhibitors (Millipore 53-913-710VL). The suspension was incubated on ice for 10 minutes. The cells were then lysed by sonication for 5 minutes, with 15-second cycles on ice. The cellular debris, containing Dps, was pelleted by centrifugation at 15,000 × g for 15 min at 4°C. The pellet was resuspended in 10 mL of 20 mM Tris at pH 8.0, 100 mM NaCl, 10 μL of 10 mg/mL DNase, and 200 μL of protease inhibitors. The resuspended pellet was incubated on ice for 10 min. The cellular debris was re-pelleted at 15,000 × g for 10 min at 4°C, and the supernatant containing the Dps protein was retained. The supernatant was concentrated to 1 mL using an Amicon centrifugal concentrator with a 10,000-MW cutoff (Millipore UFC801024). The supernatant was collected and applied to a GE MonoQ anion-exchange column, which was equilibrated in 20 mM Tris buffer at pH 8.0 and 100 mM NaCl. A 1M NaCl solution was used to elute the protein from the column. Dps was recovered from the first four column fractions. These fractions were concentrated to 0.5 mL using an Amicon 10,000-MW cut-off centrifugal concentrator. This concentrated sample was applied to a GE Superdex 200 10/30 size exclusion column equilibrated with 20 mM Tris at pH 7.0 and 100 mM NaCl. The fractions containing purified Dps were collected, and a sample was analyzed by SDS-PAGE followed by Coomassie staining. The resulting fractions containing Dps were pooled and dialyzed in buffers containing 20 mM Tris and 100 mM NaCl at pH levels of 3.0, 5.0, 7.0, 9.0, or 11.0 for use in further experiments.

### Dps-DNA Binding Gels

To determine the binding capacity of Dps across different pH values, a series of binding assays were performed. In each case, 100 ng of DNA [37] (448 bp fragmented DNA amplified via plasmids) at a concentration of 1000 ng/mL was added to a quantity of Dps, quantities ranged from 1000 to 0 ng, or in the case of pH 11, from 3500 to 0 ng (see Table 4 for further details). Buffers containing 200 mM Tris and 1 M NaCl, and pH adjusted to 3, 5, 7, 9, and 11, were used throughout the experiments when working with the Dps-DNA aliquots. Reactions were incubated at room temperature (25°C) before the addition of 2 μg of loading dye and then centrifuged at 13,000 RPM for 30 seconds. These reactions were then loaded onto a 0.5% agarose gel and run at 98 V for 60 minutes.

**Table 4.**
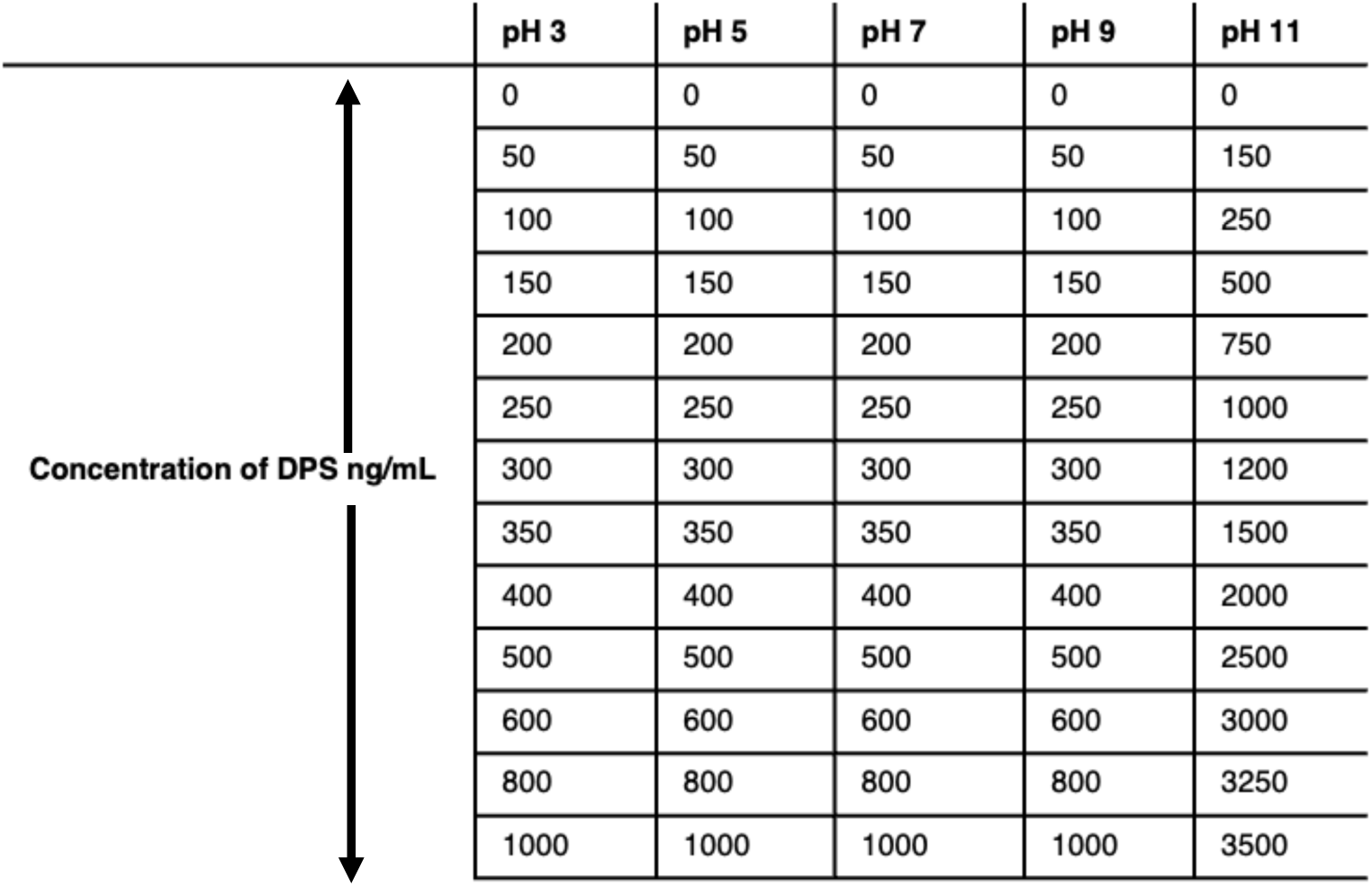
Dps concentration ranges used for DNA-binding assays across pH 3 to 11. Concentrations of purified Dps were applied in gel-based DNA-binding assays under each pH condition. For each pH, Dps was titrated over the indicated range to achieve comparable coverage of the binding curves, with the upper concentration limit adjusted to account for reduced binding affinity at higher pH values (see Table 3 and Figure 5).

### Analysis of half-maximal binding (apparent EC_50_)

Gels were imaged and band intensities quantified in ImageJ using identical rectangular ROIs for all lanes and a single background ROI per gel. We measured the free-DNA band in each lane. For each gel, a “bound like” signal was obtained from the loss of free DNA:

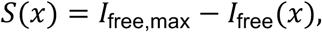

Where 𝐼_free_(𝑥) is the background-corrected integrated density of the free-DNA band at Dps concentration x, and 𝐼_free,max_ is the most significant free-DNA signal on that gel (i.e., the lane with the least binding). To place gels on the same scale, 𝑆(𝑥) was normalized to the near-saturating lane on that gel (the 1000 nM Dps lane):

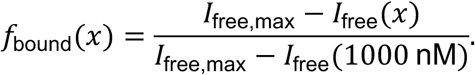

These 𝑓_bound_(𝑥) values (0–1, with 1 ≈ near saturation) were plotted versus Dps concentration. Replicates were combined by averaging 𝑓_bound_ at each concentration; per-concentration variability is shown as error bars (standard deviation). Gels were pre-screened using predefined QC criteria (monotonic trend, near-saturation at the top dose, and uniform loading/background). For each pH, three QC-passing gels were analyzed. For comparison across conditions, all five pH datasets were overlaid in a single plot of 𝑓_bound_ versus Dps concentration.

### Parametric binding-curve fitting (variable-slope 4-parameter model; performed in GraphPad Prism 10)

Each pH dataset was fit separately to the four-parameter Hill equation (also known as the ‘*[agonist] vs. response — variable slope’* model in Prism):

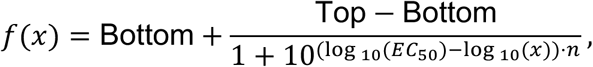

Where *Top* and *Bottom* correspond to the upper and lower plateaus, 𝐸𝐶_50_ is the apparent half-maximal concentration, and 𝑛 is the Hill slope (sensitivity parameter). Fits were performed by least-squares minimization with weighting according to replicate scatter (Prism option: *“Account for the N and scatter among replicates (from SD)”*). The *Bottom* parameter was constrained to 0, whereas *Top* was left unconstrained and allowed to vary freely. For each dataset (pH 3, 5, 7, 9, and 11), Prism reported best-fit values for *Top* (𝐵_max_ 𝐸𝐶_50_, and 𝑛 with associated standard errors and 95 % confidence intervals (profile likelihood). Goodness of fit was assessed by −𝑅^2^ and an inspection of residuals.

### Dps-DNA incubation for cryo-EM

10 µL of Dps at a concentration of 2.5 mg/mL was added to 1.5 µL of DI water and 1 µL of 200 mM Tris (1 M NaCl), whose pH was adjusted to 3, 5, 7, 9, or 11 using HCl or NaOH. 12.5 µL of 2.0 mg/mL DNA was added to the solution. This reaction was then incubated for 1 hour at room temperature.

### Cryo-EM sample preparation and data collection

For each data collection of the Dps protein (pH 3, 5, 7, 9, 11), 4 μL of sample was pipetted onto glow-discharged 300 mesh copper R1.2/1.3 Quantifoil grids. For the Dps-DNA pH 3, 7, and 11 samples, 300-mesh copper R1.2/1.3 + 2-nm continuous carbon Quantifoil grids were used.

A Vitrobot Mark IV (ThermoFisher Scientific), allowed for grid blotting for a time of 5s with 597 Whatman filter paper, and a blot force of +1 that corresponds to a relative offset from the instrument’s calibrated “zero-force” position, within a relative humidity environment of 100% and a temperature of 21 °C. These samples were then subsequently plunge-frozen into liquid ethane. Grids were stored under liquid nitrogen and then were atlased for subsequent data collection sessions.

Each high-resolution data collection was performed on a Titan Krios G3i FEG-TEM (Thermo Fisher Scientific) operating at 300 kV. Images were recorded using a Gatan K3 direct electron detector and a BioQuantum energy filter set at 20 eV. Correlated double sampling (CDS) was used to collect dose-fractioned micrographs. For each of the SPA data collections (Dps pH 3, 5, 7, 9, 11), a defocus range of -0.5 to -2.5 μm with increments of 0.25 μm was used. Data were recorded at a calibrated magnification of 105,000× (relating to a pixel size value of 0.083 Å) using the EPU software package (Thermo Fisher Scientific). Movies were recorded at a dose of 0.45 e^-^/Å²/s at 45 frames/s with a 4.25 s exposure, resulting in an accumulated total dose of 45 e^-^/Å². A total of 9,103 (pH 3), 7,613 (pH 5), 8,400 (pH 7), 7,171 (pH 9), and 7,529 (pH 11) movies were collected.

Tilt series were collected in a dose-symmetric fashion for each STA data collection (Dps at pH 3, 7, and 11). A tilt range of -57° to +57°, with increments of 3°, was applied, along with a defocus range of -3 to -5 μm, with increments of 0.5 μm. Data were recorded at a calibrated magnification of 81000× (relating to a pixel size value of 1.081 Å) using the EPU software package (Thermo Fisher Scientific). Movies were recorded at a dose of 2.62 e^-^/Å²/s at 45 frames/s with a 0.7s exposure, resulting in an accumulated total dose of 71.76 e^-^/Å². A total of 12, 52, and 12 tomograms were collected for pH 3, 7, and 11, respectively.

### Single particle cryo-EM data processing

Data processing for all SPA datasets was performed using CryoSPARC v4.6.2 [50]. The same pipeline was applied to each of these five datasets and progressed as follows: Firstly patch motion correction and CTF estimations were performed for the dataset movies, before a manual curate exposure job to remove movies deemed to have CTF fit and motion distance values larger than the consensus for the datasets (based on visual assessment of the graphical interfaces specific to each dataset).

Manual picking was then performed on approximately 10 micrographs of different defocus values. Particular attention was paid to manually picking a large number of particles visible to the human eye on each micrograph (>50 per micrograph) to maximize the capabilities of the downstream autopicking steps. 2D classification was then performed on the manually selected particles, and the best classes were used to train an initial Topaz model. This model was then used to select and extract particles from the initial 10 micrographs in a feedback loop. Following a second round of 2D classification and class selection, a new model was created from this second collection of particles. The second model was then used to select and extract particles from the entire dataset of curated movies. The initial batch of Topaz [51], auto-picked particles underwent multiple rounds of 2D classification to remove false positives and low-quality particles. From the initial quantities of auto-picked particles, the total number of good particles for the datasets was determined to be: 3,436,483 (pH 3), 4,619,982 (pH 5), 1,191,027 (pH 7), 3,087,277 (pH 9), and 842,944 (pH 11).

For each dataset, an ab initio job was generated and used as the input volume to create homo-refine jobs with both C1 and T symmetry imposed. The C1 volume functioned merely as a test to visually confirm that T was the correct order of symmetry, with no further processing being performed. The T-symmetry was subjected to subsequent iterative rounds of global CTF, local CTF, and local motion corrections, followed by homo-refinements to monitor the resulting resolution. When no further improvements were observed, reference-based motion corrections were performed, followed by another round of global and local CTF corrections. Finally, a hetero-refinement job comprising 3 classes was performed, followed by a final round of global and local CTF corrections. The resulting final particle count values attributed to 3 372 469, 4 281 527, 1 184 770, 2 974 389, 790 233 for pH values 1, 3, 5, 7, 9, and 11, respectively and were output as homo-refine jobs with final resolution values of 1.72 Å, 1.76 Å, 1.86 Å, 1.72 Å, and 1.81 Å. Supplementary Figure 11 details the Cryo-EM methodology workflow used to solve the SPA structures within this paper.

### Cryo-ET data processing and sub-tomogram averaging (STA)

Due to the lack of fiducial markers within the tilt series collections, tomograms were automatically aligned using AreTomo [40] with a VolZ value of 2460, an AlignZ value of 820, and an initial bin-factor of 6. These tomograms were then ‘corrected’ via the IsoNet [41] workflow to account for the missing wedge artifact and to increase contrast for particle picking purposes. Within the GUI workflow, default values were primarily used, with the only manually entered values being a ‘Defocus Value’ of 40000 Å, an ‘SSNR fall-off’ of 0.5, and a ‘Deconvolution Strength’ of 0.5. The extraction was run with a total value of 300 for ‘NumberSubTomo’. The extracted sub-tomograms were subsequently used to train a model (prediction) through the neural network in IsoNet [41], with 8 iterations of 32 epochs. The denoised and missing wedge corrected tomograms were ultimately then passed through Dragonfly [42], to allow for the successful segmentation of the DPS molecules through model generation for each of the pH data sets. In order to measure the approximate size of the pH 3 and pH 7 Dps-DNA structures, coordinate points were manually added to the tomograms using 3dMod [52] to calculate the distances.

### Tomogram Segmentation

Segmentation was performed in Dragonfly (Object Research Systems; v2025.1 build 14087) following Heebner et al [53]: multi-class annotations on ∼50 slices for the tomogram were manually drawn (classes: Dps, DNA, background). Following this, a 2.5D U-Net model was trained with a depth of 5, a stride ratio of 0.25, a patch size of 64, and 100 epochs, using basic augmentations (flips/rotations). The trained network was applied across the full volume; outputs were then cleaned in Dragonfly (morphological operations and manual edits), exported as binary TIFFs, and visualized in ChimeraX [54].

### Segmentation Visualization

The binary TIFF files of each segmented class were loaded into ChimeraX [54], along with the IsoNet corrected tomogram [41]. Pixel size values for the segmentation classes were corrected using the ChimeraX command line, and manual cleaning was done for each surface view using the ‘Eraser tool’. 3DMod [52] was then used to create the videos (Supplementary videos 4-6).

### Model building and validation

Alphafold 3 [31] predictions of our full-length His-tag Dps sequence (Supplementary Figure 10b) were generated as a monomer, and the highest-scoring prediction was docked into the volume of the 1.86 Å pH 7 electron density map using the ChimeraX [54] fit-to-map function. COOT [32], was used to rebuild and refine the model, which was then subjected to validation first in ISOLDE [55], and subsequently to a round of real-space refinement in Phenix [56]. ChimeraX [54] was then used to fit copies of this refined subunit, ultimately resulting in a 12mer of the 12-169AA DPS protein. This dodecamer was then validated within Phenix [56] through a final round of real-space refinement.

Iron ions were modeled at the dimer interfaces of the dodecamer, where strong residual density was observed in all twelve sites. Each Fe atom was positioned at the center of its corresponding coordination density and refined isotropically without explicit occupancy constraints. Based on map features and interatomic distances, potential coordinating residues (Asp78, Glu78, His51) were inspected in COOT [32], but no fixed LINK or metal–ligand restraints were enforced; only interactions supported by both apparent density and physically reasonable bond lengths were retained.

Local and global model–map correlation was evaluated using the Model–Map Q-score tool in ChimeraX [54]. Q-scores were calculated for the final refined model with the following parameters: points per shell = 8, shell radius step = 0.1 Å, maximum shell radius = 2.0 Å, and reference Gaussian sigma = 0.6. Mean Q-scores were determined for all atoms and, separately, for backbone and side-chain atoms to assess per-residue map-model agreement across the pH series.

All five pH maps (pH 3, 5, 7, 9, and 11) share a correlation value greater than 99% as calculated by the ‘Fit map’ program in ChimeraX [54]. Therefore, the pH 7 model was docked into each pH value electron density map. The resulting Phenix [56] validation reports confirmed that in each case, the model fits within the density maps for every pH value.

### Electrostatics

To visualize the electrostatics for DPS at various pH values, the online APBS PDB2PQR software suite was utilized (https://server.poissonboltzmann.org) [36]. pH values were set manually, and runs were performed with ‘AMBER’ force fields, ensuring that the chain IDs were retained in the PQR files. The resulting PQR files were then visualized alongside the initial PDB files to generate the images shown in Figure 3 using ChimeraX [54].

## Acknowledgements

This work was supported in part by the University of Wisconsin, Madison, the Department of Biochemistry at the University of Wisconsin, Madison, and public health service grants R01 GM104540 and U24 GM139168 to E.R.W. and F32 GM143854 to D.P. from the NIH. This work was supported in part by the U.S. Department of Energy, Office of Science, Office of Biological and Environmental Research under Award Numbers DE-SC0018409. We are grateful for the use of facilities and instrumentation at the Cryo-EM Research Center in the Department of Biochemistry at the University of Wisconsin, Madison. We are thankful for the computational resources supplied through the SBGrid Consortium ^1^.

## Data deposition

Atomic models have been deposited in the Protein Data Bank under accession XXX for Dps. Cryo-EM maps have been deposited in the Electron Microscopy Data Bank under accessions XXX for the Dps single particle cryo-EM maps.

## Supplementary Figures and Videos

**Supplementary Figure 1.**
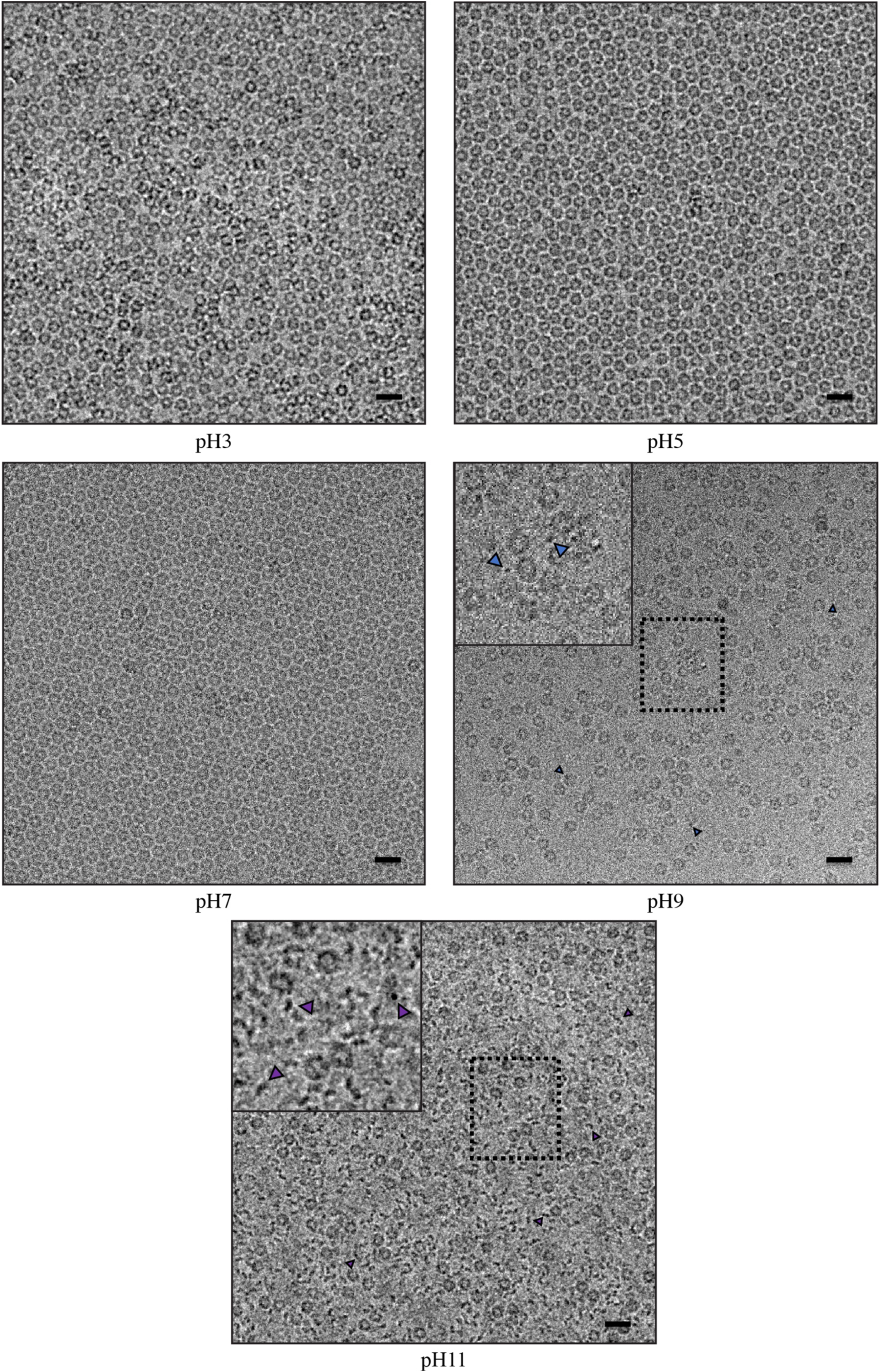
Raw micrographs of Dps SPA data collections. Raw micrograph examples of the 5 SPA data collections taken of Dps at pH values 3 (top left), 5 (top right), 7 (middle left), 9 (middle right), and 11 (bottom). For pH 9 and 11, close-ups are shown to visualize the non-uniform material also observed in the micrographs, with blue and purple arrows pointing to it in each image, respectively. Scale bar 200 Å.

**Supplementary Figure 2.**
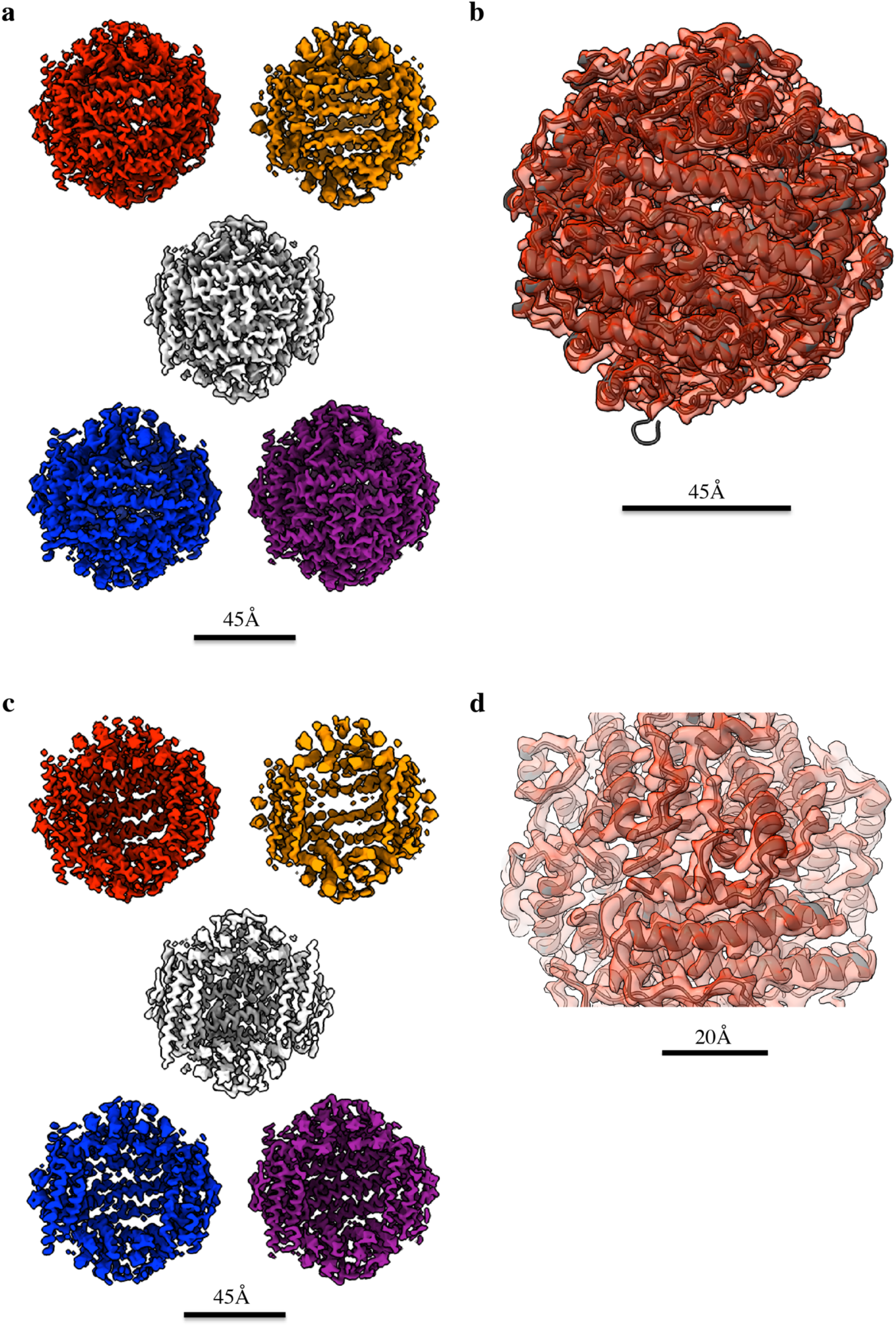
C1 reconstruction of Dps across pH conditions. (**a**) Single-particle reconstructions of Dps generated without imposed symmetry (C1) at pH 3 (red), pH 5 (orange), pH 7 (white), pH 9 (blue), and pH 11 (purple). Despite the absence of symmetry constraints, each reconstruction displays the characteristic tetrahedral (T) symmetry of the Dps dodecamer. (**b**) The X-ray crystallographic model of *E. coli* Dps (PDB code: 1DPS) fitted into the pH 3 C1 density map, demonstrating excellent correspondence between the cryo-EM and crystallographic structures. (**c**) Cross-sectional views of the same maps shown in (**a**), highlighting the internal cavity and consistent subunit packing across pH values. (**d**) Close-up of the fit shown in (**b**), illustrating the high-resolution detail and clear correspondence of secondary-structure elements. Scale bars are 45 Å (**a**, **c**) and 20 Å (**d**).

**Supplementary Figure 3.**
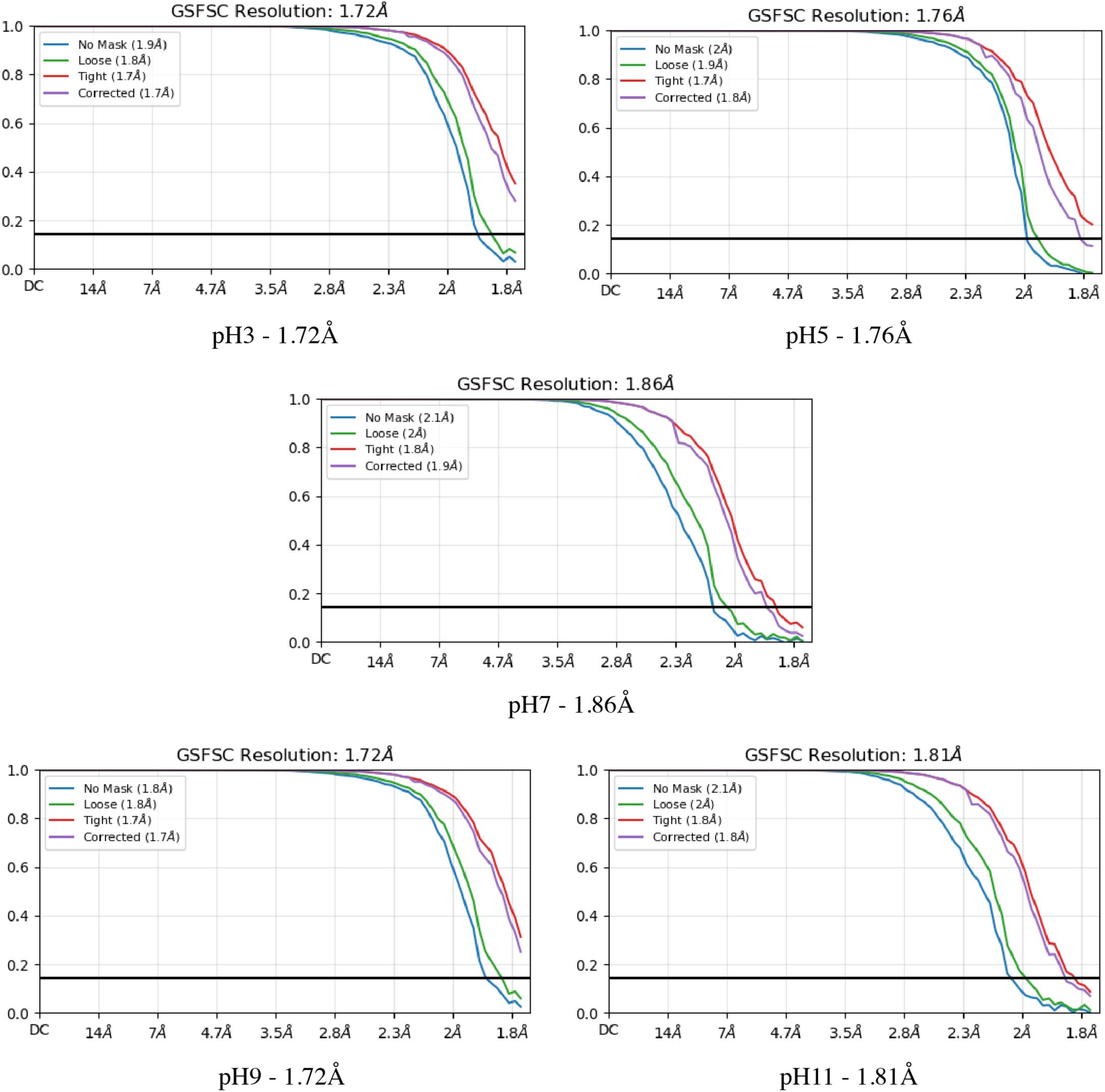
Fourier shell correlation (FSC) analysis of Dps reconstructions across pH conditions. Gold-standard FSC (GSFSC) curves for Dps at pH 3, 5, 7, 9, and 11. Curves are shown blue (no mask), green (loose mask), red (tight mask), and purple (mask-corrected FSC). The black horizontal line marks the 0.143 resolution criterion. Global resolutions (at FSC = 0.143) are 1.72 Å, 1.76 Å, 1.86 Å, 1.72 Å, and 1.81 Å for pH 3, 5, 7, 9, and 11, respectively, indicating consistently high map quality across the pH series.

**Supplementary Figure 4.**
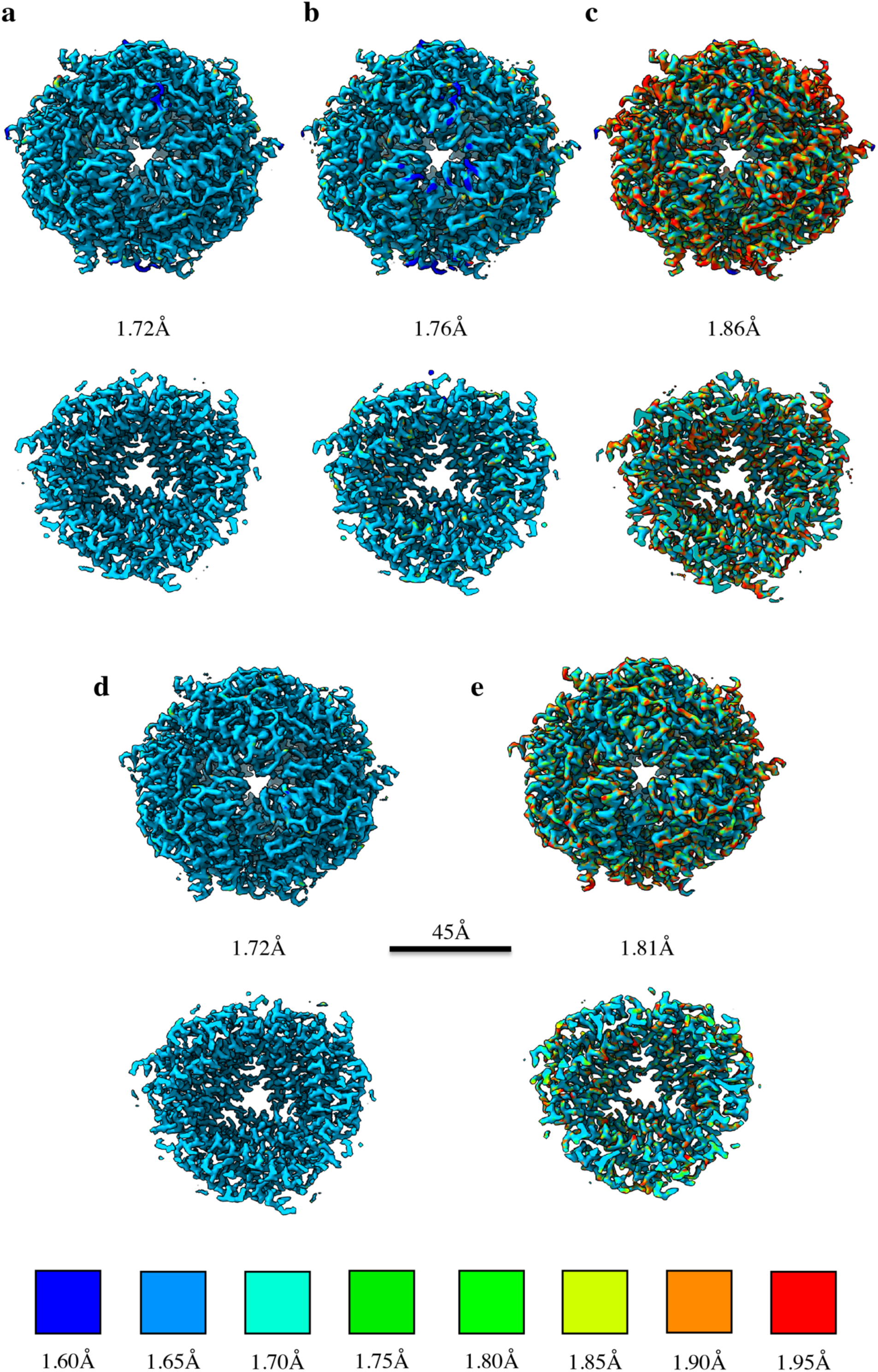
Local resolution estimation of Dps across pH conditions. Local resolution maps of the single-particle reconstructions of Dps at pH 3 (**a**), pH 5 (**b**), pH 7 (**c**), pH 9 (**d**), and pH 11 (**e**). Each map is coloured according to the local resolution scale shown below (1.60–1.95 Å), as estimated using the local resolution tool in CryoSPARC. For each reconstruction, the corresponding global resolution value determined at FSC = 0.143 is indicated beneath the map. All five datasets exhibit high overall and local resolution, with the core regions of the dodecamer achieving near-atomic detail (∼1.7 Å) and slightly lower resolution at the periphery, due to greater side-chain and terminus flexibility. Scale bar is 45 Å.

**Supplementary Figure 5.**
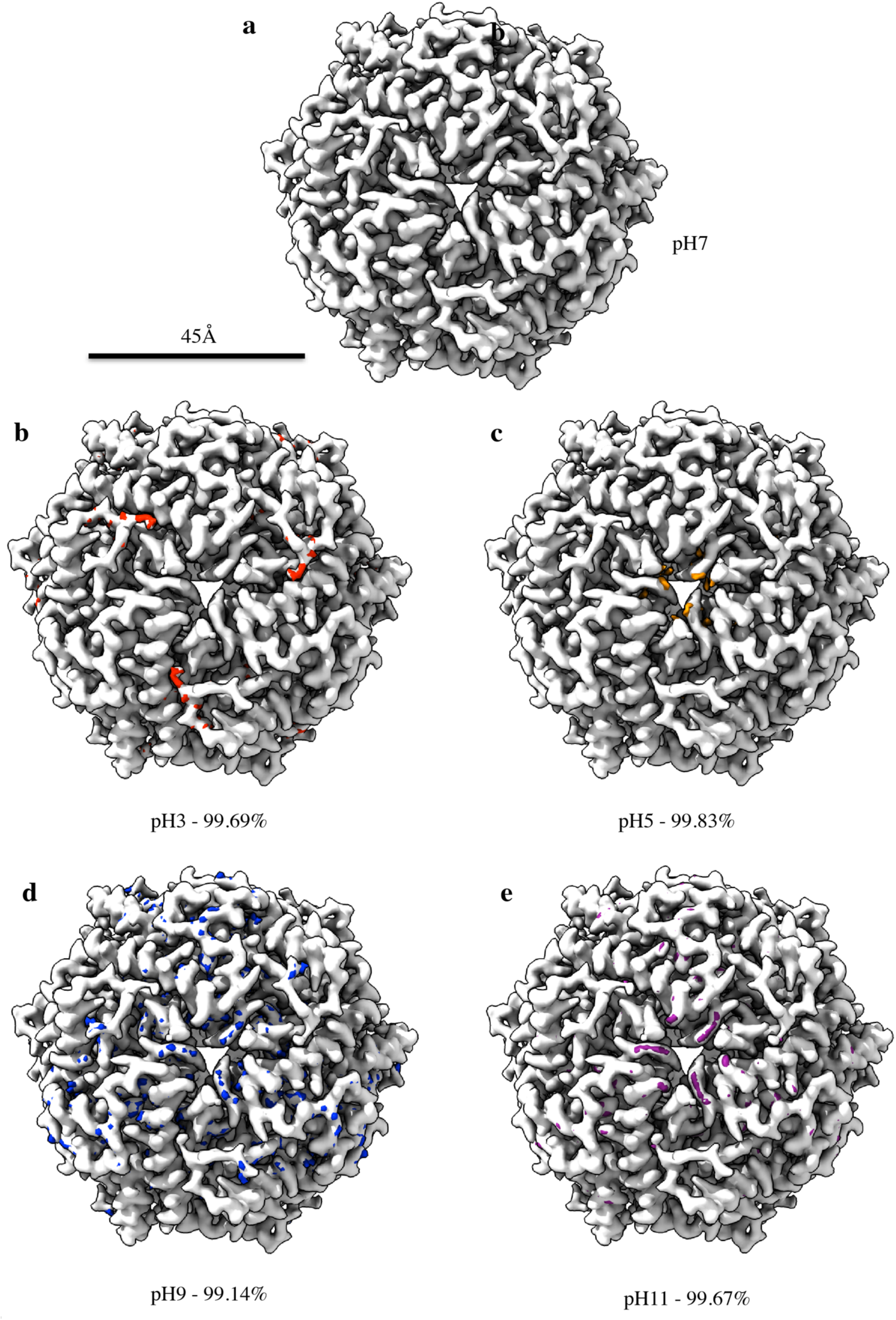
Dps density difference maps. (**a**) Electron-density map of the Dps dodecamer at pH 7, shown in white, used as the reference for comparison. (**b–e**) Difference maps showing regions of additional density relative to the pH 7 structure at pH 3 (red), pH 5 (orange), pH 9 (blue), and pH 11 (purple). These maps were generated by subtracting the pH 7 density from each respective reconstruction, thereby revealing pH-specific deviations in local conformation and side-chain positioning. The overall similarity of the maps (cross-correlation > 99 %) indicates that the global architecture of Dps remains consistent across all conditions. At the same time, the localized density variations likely correspond to subtle rearrangements of flexible termini, surface loops, or solvent-exposed residues influenced by protonation state. Scale bar is 45 Å.

**Supplementary Figure 6.**
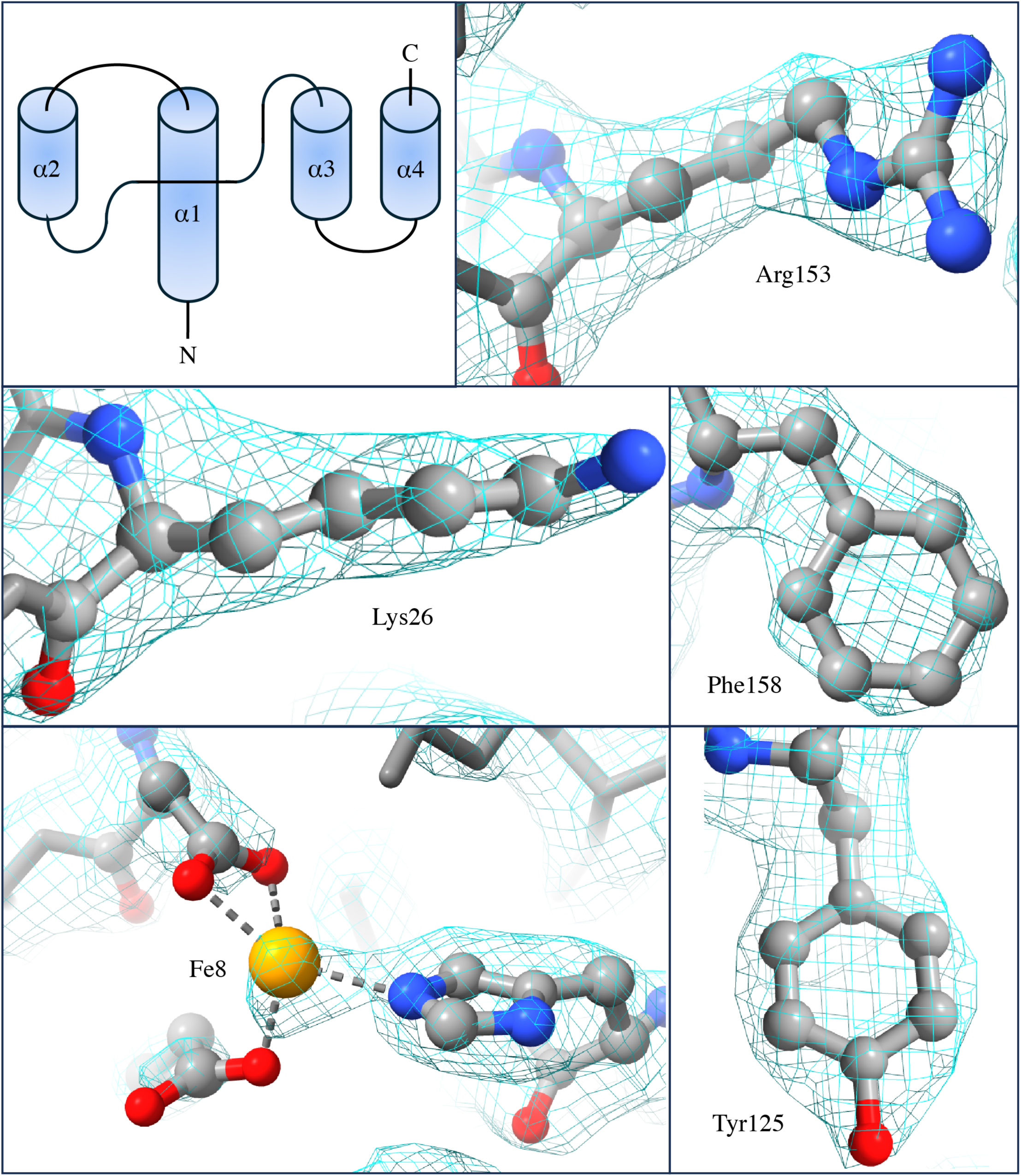
Topology and Residue fitting. Schematic topology of the Dps monomer showing its four α-helical bundles (α1–α4). Close-up panels display representative residues fitted into the 1.72 Å pH 3 electron-density map, illustrating the quality of local density and atomic detail: Arg153 (top right), Lys26 (middle left), Phe158 (middle right), and Tyr125 (bottom right). The Fe-binding site (bottom left) shows apparent density for the coordinated Fe ion and surrounding carboxylate and imidazole ligands.

**Supplementary Figure 7.**
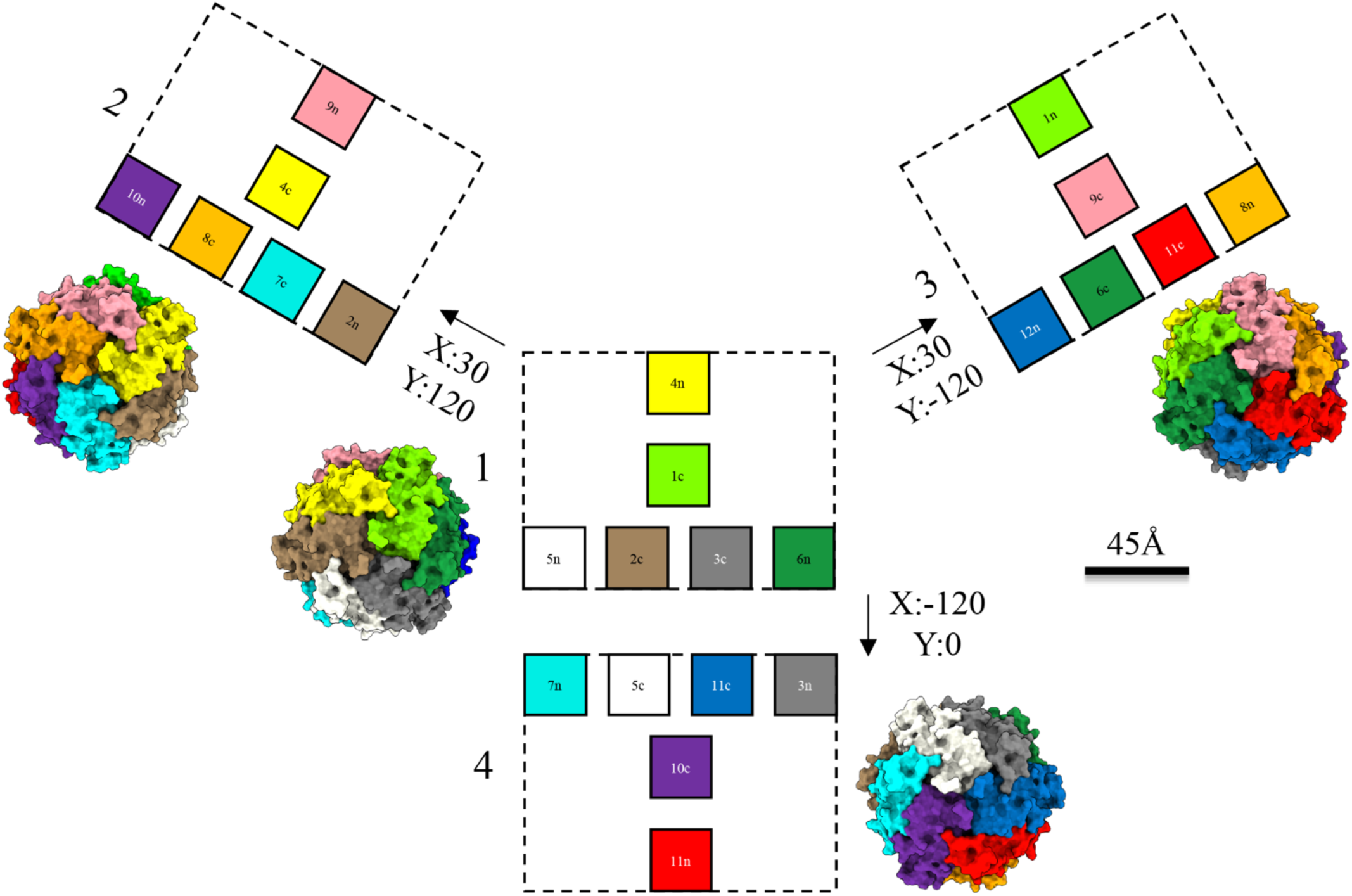
Dps dodecamer topology and trimeric organization. Schematic and three-dimensional representations of the Dps dodecamer showing how the twelve monomers are arranged into four trimeric surface units. Each dashed rectangle (labeled 1–4) depicts a trimeric assembly, viewed in two dimensions to illustrate the spatial relationships between the N- and C-terminal regions of different monomers. Within each rectangle, the N-termini of three subunits surround the C-termini of three neighboring subunits, forming combined pseudo-hexameric clusters. As an example, monomer 5 has its N-terminus (*5n*, white box) located within rectangle 1 and its C-terminus (*5c*, white box) in rectangle 4, demonstrating how alternating N- and C-terminal interactions interconnect across the dodecamer surface. This reciprocal pattern is consistent across all 12 subunits, yielding 4 N/C-terminal trimer pairs that together form the complete dodecameric architecture. The corresponding 3D surface renderings adjacent to each rectangle show the actual Dps density and surface features for the respective trimeric units. Rotational transformations along the X and Y axes (indicated by arrows) illustrate the geometric relationship between each trimeric group.

**Supplementary Figure 8.**
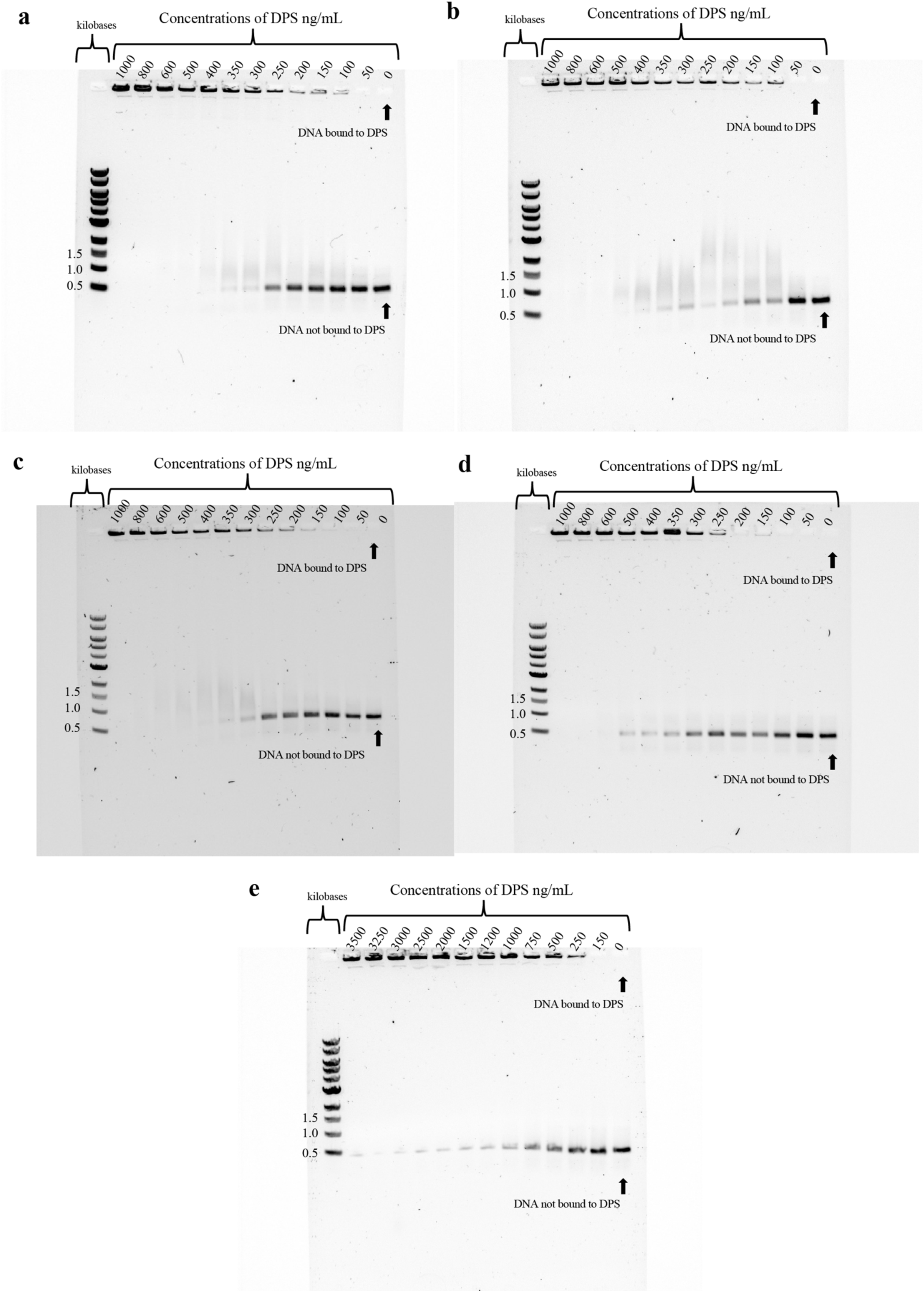
Electrophoretic mobility shift assays showing pH-dependent DNA binding by Dps. Agarose gels with DNA visualized by ethidium bromide staining, showing how increasing Dps concentration alters DNA mobility at **(a)** pH 3, **(b)** pH 5, **(c)** pH 7, **(d)** pH 9, and **(e)** pH 11. Each lane contains an identical amount of DNA with increasing Dps concentrations (0–1000 ng mL⁻¹ in **a–d**; 0–3500 ng mL⁻¹ in **e**, as indicated). Upon binding, Dps forms large DNA–protein complexes that remain near the well, reducing the intensity of the free-DNA band migrating toward the bottom of the gel. At acidic/neutral pH (**a–c**), even low Dps concentrations cause nearly complete retention of DNA, whereas higher concentrations are required for binding at neutral and alkaline pH (**d-e**). These results show that Dps–DNA complex formation is strongly pH-dependent, with maximal binding under mildly acidic conditions.

**Supplementary Figure 9.**
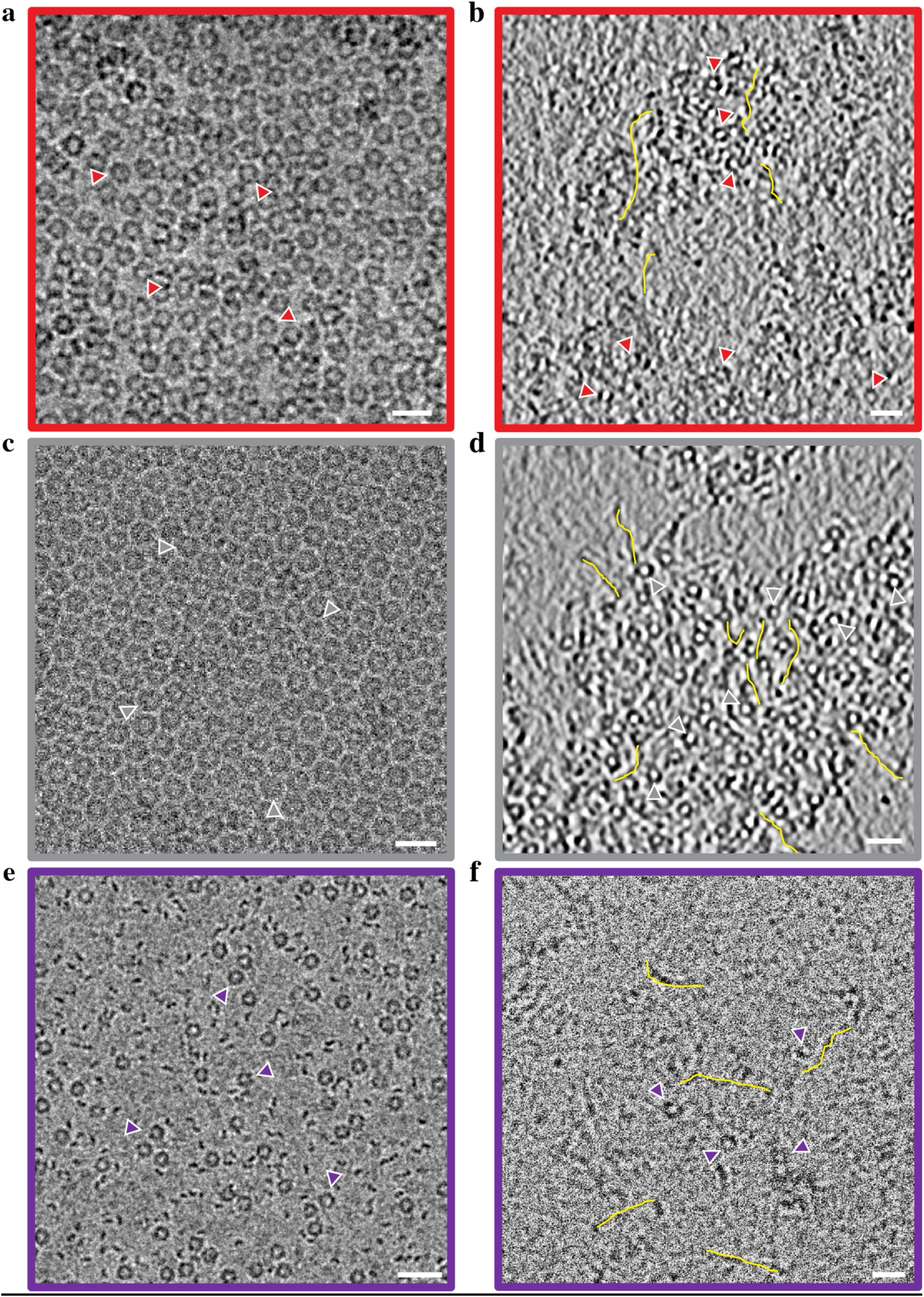
Dps-DNA CryoET. **(a**) SPA micrograph of Dps at pH 3, visualizing its ability to form 2D lattices in the absence of DNA. Red arrows point to individual Dps dodecamers. (**b**) reconstructed tomogram z-slice of Dps at pH 3 in the presence of DNA, visualizing its ability to interact with both other Dps 12mers and DNA to form significant Dps-DNA globoids. For ease, yellow lines have been drawn for some DNA lengths, and red triangles indicate Dps dodecamers in proximity to the DNA. (**c**) SPA micrograph of Dps at pH 7, visualizing its ability to form 2D lattices in the absence of DNA. Grey arrows point to individual Dps dodecamers. (**d**) reconstructed tomogram z-slice of Dps at pH 7 in the presence of DNA, visualizing its ability to interact with both other Dps 12mers and DNA to form large Dps-DNA globoids. For ease, yellow lines have been drawn for some DNA lengths, and grey triangles indicate Dps dodecamers in proximity to the DNA. (**e**) SPA micrograph of Dps at pH 11, showing its lack of ability to form a 2D lattice; instead, Dps 12-mers don’t interact with each other. (**f**) reconstructed tomogram z-slice of Dps at pH 11 in the presence of DNA, visualizing poor ability to interact with each other. For ease, yellow lines have been drawn for some DNA lengths, and purple triangles indicate Dps dodecamers. Scale bars are 200 Å.

**Supplementary Figure 10.**
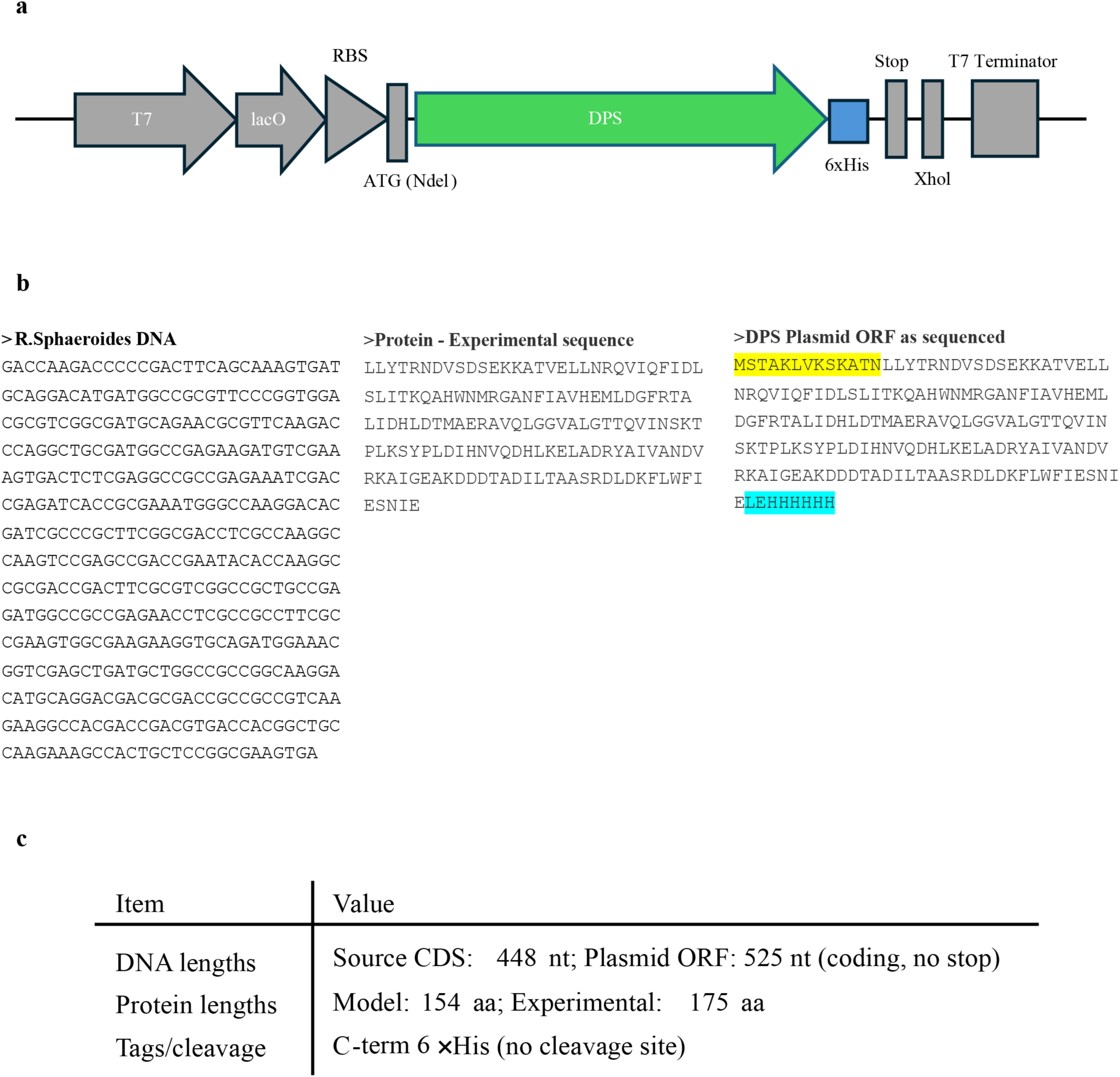
Dps construct and sequence comparison. (**a**) Simplified expression cassette from the sequenced pET plasmid showing regulatory and cloning features (T7 → lacO → RBS → NdeI–ATG → *DPS* → 6×His → Stop → T7 terminator). Only the expression region is shown (backbone elements such as ori, Amp, lacI omitted for clarity). (**b)** Comparison of the *R. sphaeroides* genomic *dps* coding sequence, the experimentally determined protein, and the plasmid open reading frame (ORF) as sequenced. Vector-derived residues (tags/linkers) are highlighted in **yellow** or **blue**. (**c**) Summary of sequence lengths, tags, and substitutions.

**Supplementary Figure 11.**
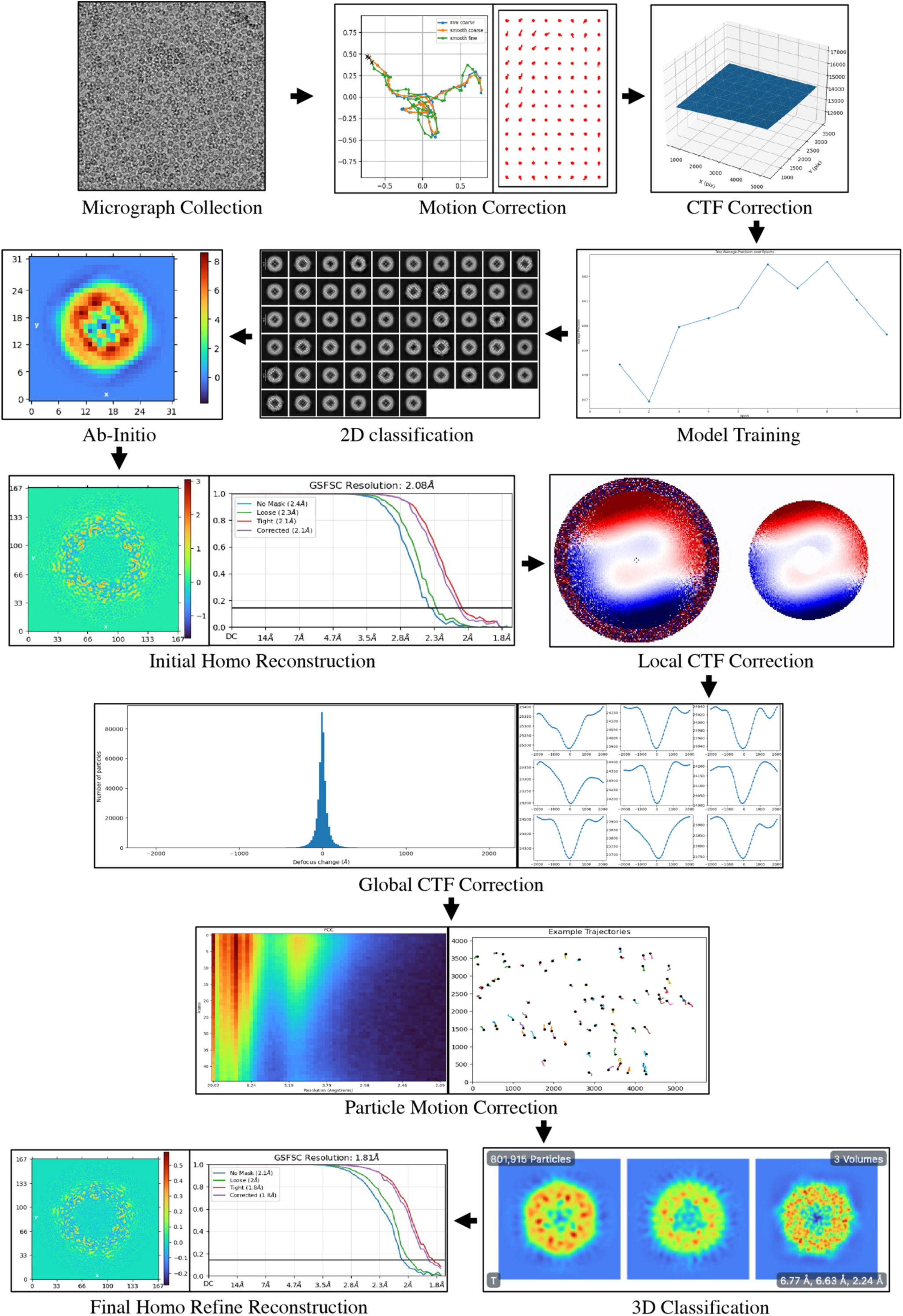
Workflow for determining the Dps structure by cryo-EM. Cryo-EM data processing pipeline used to solve the Dps structure. Following sample vitrification, high-quality transmission electron microscopy (TEM) micrographs were collected. Motion and contrast transfer function (CTF) corrections were applied to the raw micrographs. Afterward, Topaz model training in CryoSPARC was used to identify Dps particles and exclude background noise. Multiple rounds of 2D classification were performed to remove low-quality particles prior to *ab initio* reconstruction, yielding an unbiased initial 3D volume. This model was refined through homogeneous refinement with tetrahedral symmetry imposed. The reconstruction was subsequently improved through local and global CTF corrections, as well as particle motion correction. Final 3D classification served as an additional particle filter, leading to the generation of the final homogeneous refinement map at the highest resolution. This workflow was applied identically to all Dps datasets collected across the pH series.

**Supplementary video 1. Dps-DNA tomogram at pH 3.** Video of the tomographic reconstruction (Supplementary Figure 9b) of Dps-DNA interactions at pH 3.

**Supplementary video 2. Dps-DNA tomogram at pH 7.** Video of the tomographic reconstruction (Supplementary Figure 9d) of Dps-DNA interactions at pH 7.

**Supplementary video 3. Dps-DNA tomogram at pH 11.** Video of the tomographic reconstruction (Supplementary Figure 9f) of Dps-DNA interactions at pH 11.

**Supplementary video 4. Dps-DNA tomogram segmentation at pH 3.** Video of the segmentation (Figure 6a) of Dps-DNA interactions at pH 3, where Dps are shown in red and DNA in yellow.

**Supplementary video 5. Dps-DNA tomogram segmentation at pH 7.** Video of the segmentation (Figure 6b) of Dps-DNA interactions at pH7, where DPS are shown in grey and DNA in yellow.

**Supplementary video 6. Dps-DNA tomogram segmentation at pH 11.** Video of the segmentation (Figure 6c) of Dps-DNA interactions at pH11, where Dps are shown in purple and DNA in yellow.

